# RIPK3 promotes neuronal survival by suppressing excitatory neurotransmission during CNS viral infection

**DOI:** 10.1101/2024.04.26.591333

**Authors:** Irving Estevez, Benjamin D. Buckley, Nicholas Panzera, Marissa Lindman, Tsui-Wen Chou, Micheal McCourt, Brandon J. Vaglio, Colm Atkins, Bonnie L. Firestein, Brian P. Daniels

## Abstract

While recent work has identified roles for immune mediators in the regulation of neural activity, the capacity for cell intrinsic innate immune signaling within neurons to influence neurotransmission remains poorly understood. However, the existing evidence linking immune signaling with neuronal function suggests that modulation of neurotransmission may serve previously undefined roles in host protection during infection of the central nervous system. Here, we identify a specialized function for RIPK3, a kinase traditionally associated with necroptotic cell death, in preserving neuronal survival during neurotropic flavivirus infection through the suppression of excitatory neurotransmission. We show that RIPK3 coordinates transcriptomic changes in neurons that suppress neuronal glutamate signaling, thereby desensitizing neurons to excitotoxic cell death. These effects occur independently of the traditional functions of RIPK3 in promoting necroptosis and inflammatory transcription. Instead, RIPK3 promotes phosphorylation of the key neuronal regulatory kinase CaMKII, which in turn activates the transcription factor CREB to drive a neuroprotective transcriptional program and suppress deleterious glutamatergic signaling. These findings identify an unexpected function for a canonical cell death protein in promoting neuronal survival during viral infection through the modulation of neuronal activity, highlighting new mechanisms of neuroimmune crosstalk.

## Introduction

While the field of neuroimmunology has historically focused on roles for immune cells and molecules in driving the pathogenesis of neurological disorders, recent advances have refined our understanding of neuroimmune crosstalk to include indispensable roles for the immune system in nervous system development, homeostasis, neuroprotection, and repair^1-4^. Newly appreciated links between these two systems include unexpected roles for neuronal innate immune signaling in the regulation of neural activity^5,6^. Within this context, immune mediators transcend their traditional functions in inflammation and pathogen control and become key players in the modulation of neural circuits, influencing processes ranging from neurotransmission to behavior^7-9^. Defining the mechanisms and consequences of neuroimmune signaling has, therefore, become paramount to understanding the basic biology of the nervous system, with implications for the development of therapeutic strategies during disease.

Recent work has described specialized adaptations of several innate immune processes in neurons. For example, we and others have described cell death-independent functions for Receptor interacting kinase-3 (RIPK3), the canonical inducer of a form of programmed cell death termed “necroptosis”^10-12^. In the setting of neurotropic flavivirus infection, activation of RIPK3 does not engage the necroptotic executioner molecule mixed lineage kinase domain-like protein (MLKL), but rather coordinates a dramatic shift in neuronal transcription and metabolism to induce an antiviral state in the absence of cell death^13-15^. This outcome may represent an adaptive strategy to control infection without sacrificing a critical postmitotic cell type that cannot be replaced in the adult brain^16,17^. Despite these insights, the mechanisms by which RIPK3 shapes the neuronal transcriptome during infection are poorly understood, as are the ways in which engagement of this pathway influences neuronal cell biology beyond inducing antimicrobial gene expression.

Notably, aberrant neuronal activity appears to be a major driver of disease pathogenesis during viral infections of the central nervous system^18-22^. Perturbations to excitatory neurotransmission mediated by glutamate have now been linked to the neurologic damage elicited by neurotropic flaviviruses, including the major human pathogens West Nile virus (WNV), Zika virus (ZIKV), and Japanese encephalitis virus (JEV)^23-26^. Flavivirus induced-enhancement of glutamate signaling results in excitotoxicity, a pathological process triggered by the excessive activation of ionotropic glutamate receptors on neuronal cells. Excessive influx of calcium ions through the N-methyl-D-aspartate (NMDA) receptor, in particular, results in a cascade of events that culminate in neuronal damage and death^27^. While previous work has broadly identified roles for innate immune cytokines in sensitizing neurons to excitotoxicity^28,29^, the mechanisms by which viral infections shape neuronal excitability and, by extension, susceptibility to excitotoxic cell death remain poorly understood. Such insight is critically needed as modulation of neuronal activity may represent an underexplored avenue of therapeutic development for neuroinvasive flavivirus infections, which are associated with a constellation of severe neurologic syndromes and for which there are currently no disease-specific treatments^30,31^.

Here, we define an unexpected function for RIPK3 signaling in suppressing neuronal excitability during flavivirus infection, thereby promoting *survival* rather than cell death under excitotoxic conditions. Using both conditional *Ripk3* deletion and an inducible chemogenetic RIPK3 activation system, we show that RIPK3 activity in neurons drives transcriptomic changes that result in diminished sensitivity to glutamatergic stimulation, and that engagement of RIPK3 preserves neuronal viability and host survival during both flavivirus infection and sterile excitotoxic insults. Mechanistically, we show that the neuroprotective function of RIPK3 requires the activity of Ca2^+^/calmodulin-dependent protein kinase II (CaMKII), which directly associates with activated RIPK3 in protein pulldown experiments and is phosphorylated in a RIPK3-dependent manner during flavivirus infection. This RIPK3/CaMKII axis activates the transcription factor cAMP Response Element-Binding Protein (CREB), which is also required for RIPK3-mediated suppression of excitotoxicity. We show that RIPK3-dependent engagement of CREB drives the expression of a neuroprotective transcriptional program, identifying a previously unknown mechanism of RIPK3-mediated transcriptional regulation in neurons that is independent of Nuclear factor-κB (NF-κB)-dependent inflammatory gene expression or MLKL-dependent necroptosis. Together, these findings highlight mechanisms of neuroimmune crosstalk in which cell intrinsic innate immune signaling in neurons can regulate fundamental, non-immune aspects of neuronal function, including neurotransmission.

## Results

### RIPK3 is a key regulator of neurologic gene expression during flavivirus infection

To investigate roles for RIPK3 in shaping neuronal responses to flavivirus infection, we performed secondary analysis of a transcriptomic dataset previously published by our group and others^13^ in which primary cortical neuron cultures derived from *Ripk3*^−/−^ mice or wild-type (*Ripk3*^*+/+*^) littermate controls were infected with either ZIKV-MR766 or WNV-TX02 for 24h. While we had previously used this dataset to identify mediators of immunometabolic regulation in neurons, we now sought to profile the RIPK3-dependent transcriptional program more comprehensively in flavivirus-infected neurons. UMAP analysis revealed tight and distinct clustering that segregated samples by both genotype and infection status, suggesting clear RIPK3-dependent transcriptional responses to both viruses (Figure 1A). Further analysis revealed robust differential gene expression in *Ripk3*^*+/+*^ neurons in response to both ZIKV (Figure 1B) and WNV (Figure 1C) which was predominated by upregulated transcripts. Notably, *Ripk3*^−/−^ neurons exhibited hundreds of differentially expressed genes (DEGs) compared to *Ripk3*^*+/+*^ controls following infection with both viruses (Figure 1D-E). Gene ontology (GO) enrichment analysis of significant DEGs revealed overrepresentation of terms related to immune activation, especially immune cell chemotaxis, when comparing *Ripk3*^−/−^ neurons to *Ripk3*^*+/+*^ controls in both infection groups (Figure 1F-G), consistent with our previous work demonstrating a central for RIPK3 in promoting expression of chemokines and other innate immune genes in neurons during flavivirus infection^13-15^. To our surprise, however, these comparisons also revealed enrichment of GO terms associated with neurologic functions, including terms related to neurotransmission such as synaptic signaling, long-term potentiation, and neurotransmitter release (Figure 1H-I). These data suggest that RIPK3 may shape features of neuronal cell biology that extend beyond inflammatory signaling during viral infection.

**Figure 1.**
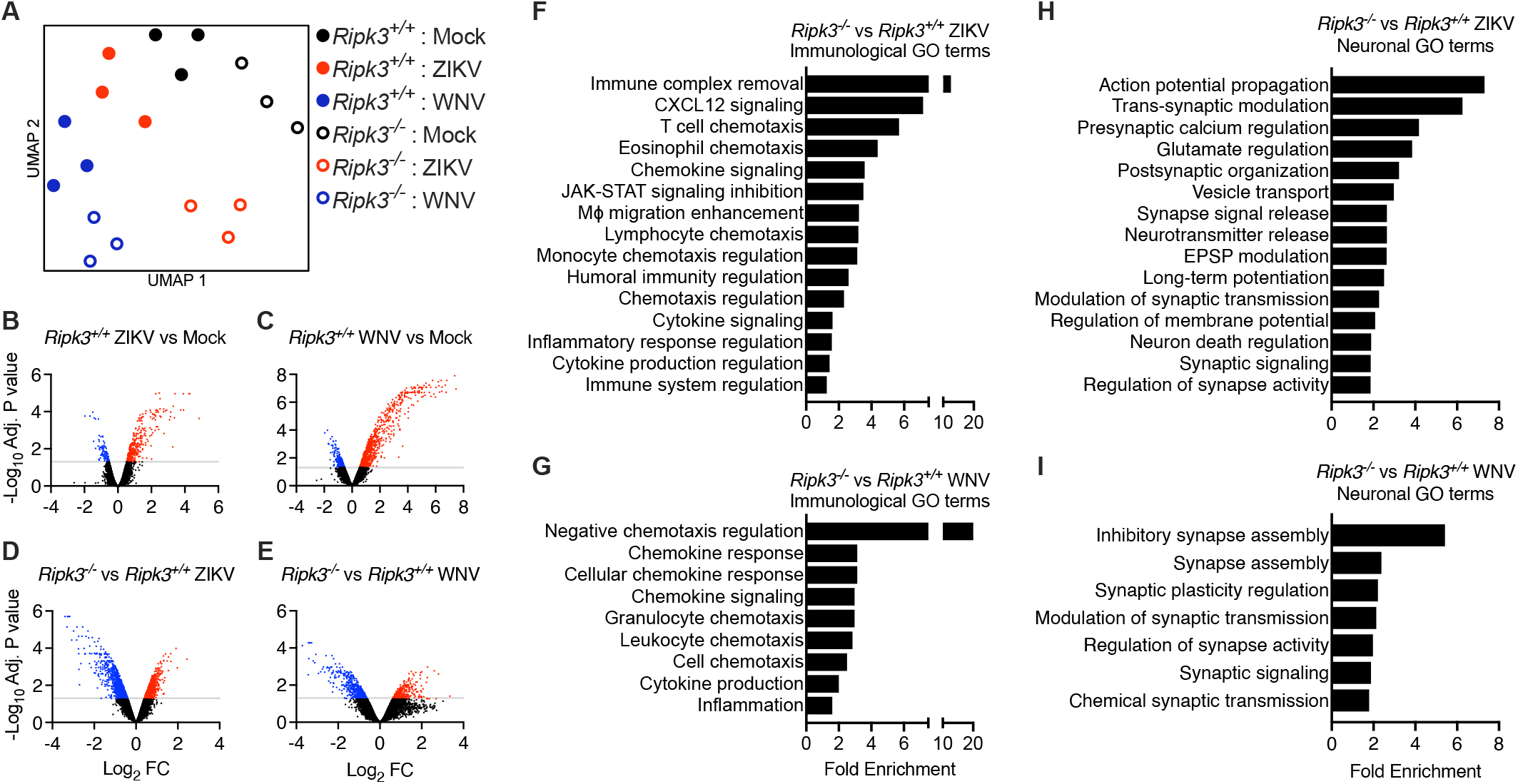
RIPK3 is a key regulator of neurologic gene expression during flavivirus infection. (A) Uniform Manifold Approximation and Projection analysis of transcriptomic data from primary cortical neurons of indicated genotypes following 24h infection with ZIKV or WNV. (B-E) Volcano plots illustrating differential expression of transcripts in neuronal cultures described in (A). Transcripts with significant changes (>1.5-fold change, adj. P < 0.05) are highlighted: downregulated transcripts are show in blue and upregulated in red. Values in (B-C) represent values in WT (*Ripk3*^*+/+*^) neurons infected with ZIKV (B) or WNV (C) compared to uninfected WT control neurons. Values in (D-E) represent values in *Ripk3*^−/−^ neuron cultures infected with ZIKV (D) or WNV (E) compared to *Ripk3*^*+/+*^ cultures infected with the same virus. (F-I) Selected overrepresented GO terms obtained from GO enrichment analysis of DEGs between *Ripk3*^−/−^ and *Ripk3*^*+/+*^ neurons infected with ZIKV or WNV, highlighting alterations in immunological (F-G) and neurological pathways (H-I). All terms were overrepresented with a FDR <0.05.

### RIPK3 signaling during flavivirus infection promotes neuronal survival rather than cell death

Recent work has implicated excitotoxicity mediated by aberrant glutamatergic neurotransmission in promoting neuronal death during flavivirus infection^20,32-34^. We thus questioned whether the RIPK3-dependent neuronal transcriptional response included genes relevant for glutamate signaling. Further analysis of genes related to glutamate receptor signaling confirmed this to be the case, with *Ripk3*^−/−^ neurons exhibiting a complex pattern of up- and down-regulated transcripts among significant DEGs for both ZIKV and WNV infection groups (Figure 2A). To better understand the implications of these findings, we performed pathogenesis studies using well established pharmacologic agents that selectively antagonize the ionotropic glutamate receptors AMPAR and NMDAR, both of which mediate excitatory neurotransmission by facilitating postsynaptic cation influx (Figure 2B). We first assessed survival in immunocompetent mice following intracranial infection with ZIKV-MR766 (Figure 2C). We observed that *Ripk3*^−/−^ mice exhibited dramatic enhancement of mortality in this model (Figure 2D), consistent with our previous work^13^. Strikingly, however, this effect could be completely ameliorated via administration of the AMPAR antagonist GYKI 52466 (GYKI) daily on days 3-7 post infection. We performed similar experiments in mice harboring neuron-specific deletion of *Ripk3* by crossing mice in which exons 2 and 3 of the endogenous *Ripk3* locus are flanked by loxP sites^35^ (*Ripk3*^fl/fl^) to a line expressing Cre-recombinase under the control of the Synapsin-1 (*Syn1*) promoter^36^. *Ripk3*^fl/fl^ *Syn1* Cre^+^ mice also exhibited significantly enhanced mortality following intracranial ZIKV or subcutaneous WNV infections compared to littermate Cre^-^ controls, and this phenotype could also be ameliorated by treatment with GYKI (Figure 2E-2F), suggesting that antagonism of excitatory glutamatergic neurotransmission through AMPAR was sufficient to rescue the enhanced mortality resulting from loss of neuronal RIPK3. We observed similar results using the NMDAR antagonist MK801, which rescued the enhanced mortality observed in both ZIKV-infected *Ripk3*^−/−^ mice (Figure 2G) and ZIKV or WNV-infected *Ripk3*^fl/fl^ *Syn1* Cre^+^ mice (Figure 2H-2I). These results further support the idea that enhanced viral pathogenesis in mice lacking neuronal RIPK3 is driven by excitotoxic glutamatergic neurotransmission.

**Figure 2.**
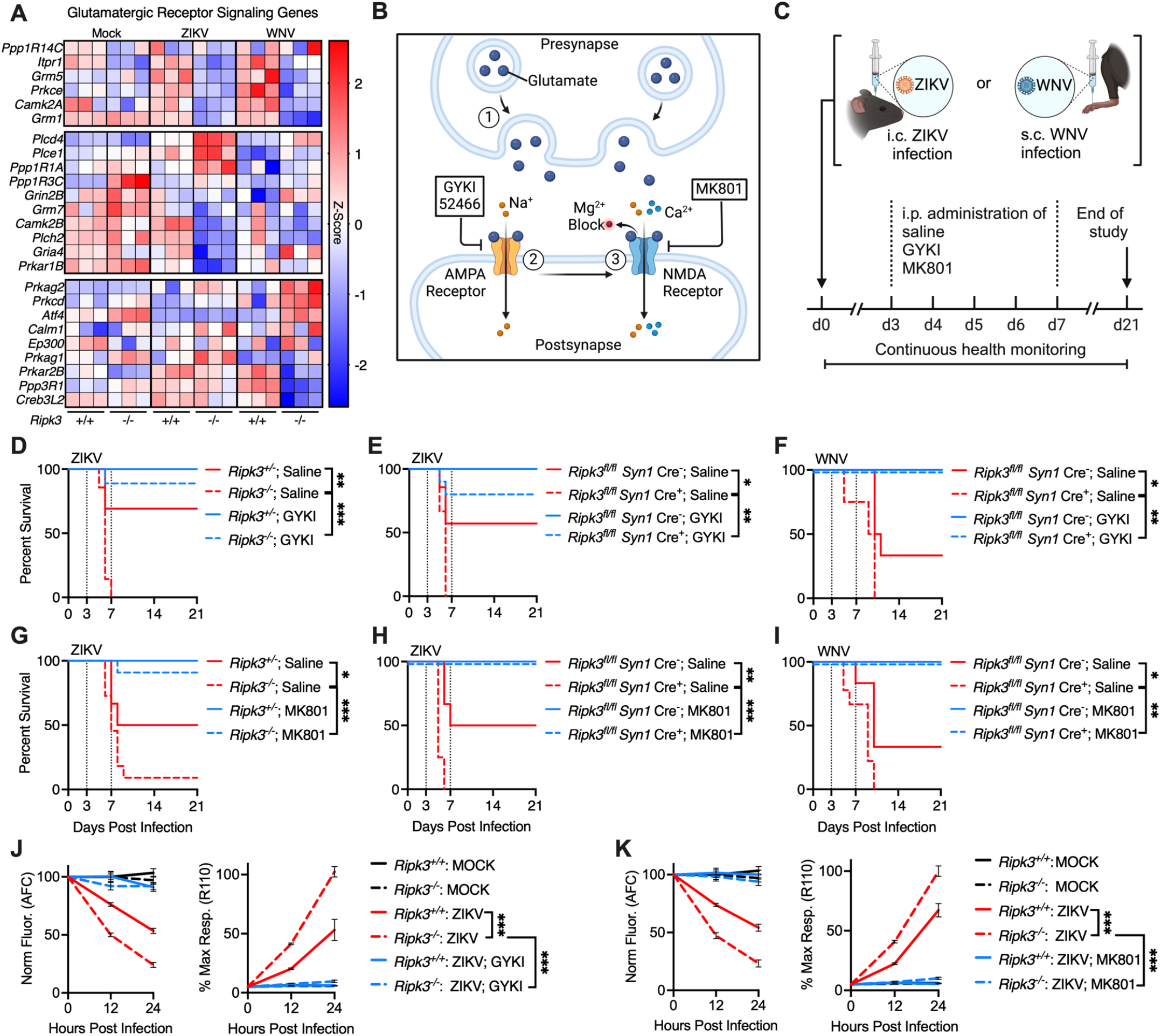
RIPK3 signaling during flavivirus infection promotes neuronal survival rather than cell death. (A) Heatmap depicting selected glutamatergic receptor signaling-associated genes derived from microarray profiling of primary cortical neurons of indicated genotypes following 24h infection with ZIKV or WNV. (B) Schematic representation of glutamatergic signaling and inhibitors used in our study. Note sequential influx of Na^+^ through AMPA receptors (blocked by GYKI-52466) and Na^+^/Ca2^+^ through NMDA receptors (blocked by MK801). (C) Schematic representation of survival studies in which mice underwent either intracranial ZIKV or subcutaneous WNV infections. Animals were treated with either saline, GYKI-52466 (1μg/g) or MK801 (0.06μg/g) on days 3-7 post infection and monitored for survival. (D-I) Survival analysis in mice of indicated genotypes following ZIKV or WNV infection with or without 5 daily treatments using GYKI-52466 (D-F), or MK801 (G-I). N = 6-13 mice per genotype ZIKV studies and N = 4-5 for WNV studies. (J-K) MultiTox cell death assays evaluating glutamatergic signaling on neuronal survival: Live (AFC) and dead (R110) protease activities were measured in primary neuron cultures from indicated genotypes infected with ZIKV and pretreated with GYKI-52466 (J) or MK801 (K). Fluorescence for AFC is normalized to mock controls, and percentage of maximum R110 response is normalized to the group exhibiting highest signal intensity at 24 hours. N = 6 independent cultures per group/condition. **p < 0.01, ***p < 0.001. Error bars represent SEM. **See also Figure S1**

To further test this idea, we assessed viability in primary neuron cultures following ZIKV-MR766 infection using the MultiTox Cytotoxicity Assay, which simultaneously measures unique protease activities associated with living and dead cells using distinct fluorogenic peptide substrates^37^. Notably, we observed that *Ripk3*^−/−^ neurons exhibited diminished live cell protease activity (AFC) and enhanced dead cell protease activity (R110) following ZIKV infection (Figure 2J), demonstrating that RIPK3 signaling promoted *survival* rather than cell death in this setting. Treatment with GYKI completely abrogated loss of cell viability in infected neurons in both assays, suggesting that the enhanced cell death occurring in the absence of RIPK3 was dependent on AMPAR-mediated glutamate signaling (Figure 2J). We observed essentially identical outcomes in experiments using the NMDAR antagonist MK801 (Figure 2K), and further confirmed these results using a traditional ATP-based viability assay (Figure S1A). We did not observe any impact of glutamate receptor inhibitors on brain viral burden *in vivo* (Figure S1B-C), nor on ZIKV replication in primary neuron cultures (Figure S1D), suggesting that the protective effect of glutamate receptor blockade was not a direct antiviral effect. We also confirmed that, while ZIKV induced enhanced glutamate release in cultured neurons (consistent with previous reports^32^), this effect did not differ between *Ripk3*^*+/+*^ and *Ripk3*^−/−^ cultures (Figure S1E), suggesting that differences in survival were not driven by differential levels of glutamate release into culture media. Furthermore, we did not observe differences between genotypes following sterile exposure to varying concentrations of exogenous glutamate or NMDA, indicating that *Ripk3* deficiency alone does not alter baseline sensitivity to glutamate- or NMDA-induced toxicity (Figure S1F-1G). Together, these data support the idea that flavivirus infection induces glutamatergic excitotoxicity and that activation of neuronal RIPK3 signaling suppresses this effect.

### RIPK3 modulates neuronal excitability and promotes neuroprotection during excitotoxicity

Previous work has shown that glutamate-induced neuronal excitotoxicity is initiated by an overabundance of intracellular Ca^2+^ ions, which initiate a variety of intracellular signals that culminate in cell death^38-40^. More recently, increased cytoplasmic Ca^2+^ influx via NMDARs has also been shown to drive neuronal cell death during viral infection^32^. We therefore questioned whether prophylactic engagement of RIPK3 could suppress neuronal cell death in the setting of excitotoxic stimulation. To this end, we generated primary cortical neuron cultures from transgenic mice expressing a chemogenetically activatable form of RIPK3 (RIPK3-2xFV)^14^ under the control of the *Nestin* promoter (*Ripk3*-2xFV^fl/fl^ *Nestin* Cre^+^). RIPK3-2xFV proteins contain tandem FKBP^F36V^ domains that drive forced oligomerization and activation of RIPK3 following exposure to the dimerization drug B/B homodimerizer (B/B) (Figure 3A). Using this system, we can thus induce sterile activation of RIPK3 in the absence of any exogenous stimulus. We validated the expression of RIPK3-2xFV through immunocytochemical staining of the reporter mCherry (Figure 3B). We also confirmed that B/B treatment induced the expression of *Cxcl10* (Figure S2A), which we and others have shown to be strongly induced by RIPK3 in a variety of settings^13,14,41^. We first pretreated RIPK3-2xFV-expressing neuron cultures with B/B for 24h, followed by infection with ZIKV-MR766 and assessment of cell viability using the MultiTox Cytotoxicity assay. B/B pretreatment robustly suppressed neuronal cell death induced by ZIKV infection, further supporting a pro-survival function of this molecule in neurons (Figure 3C). Remarkably, B/B pretreatment also protected neurons from an excitotoxic dose of exogenous NMDA, confirming that activated RIPK3 can directly impact the cellular outcomes of glutamatergic receptor activation (Figure 3D). We repeated these experiments using an alternative ATP-based viability assay, confirming that B/B pretreatment was neuroprotective against both ZIKV and NMDA insult in *Ripk3*-2xFV^fl/fl^ *Nestin* Cre^+^ neurons, but not in Cre^-^ cultures which lack RIPK3-2xFV transgene expression (Figure S2B-C).

**Figure 3.**
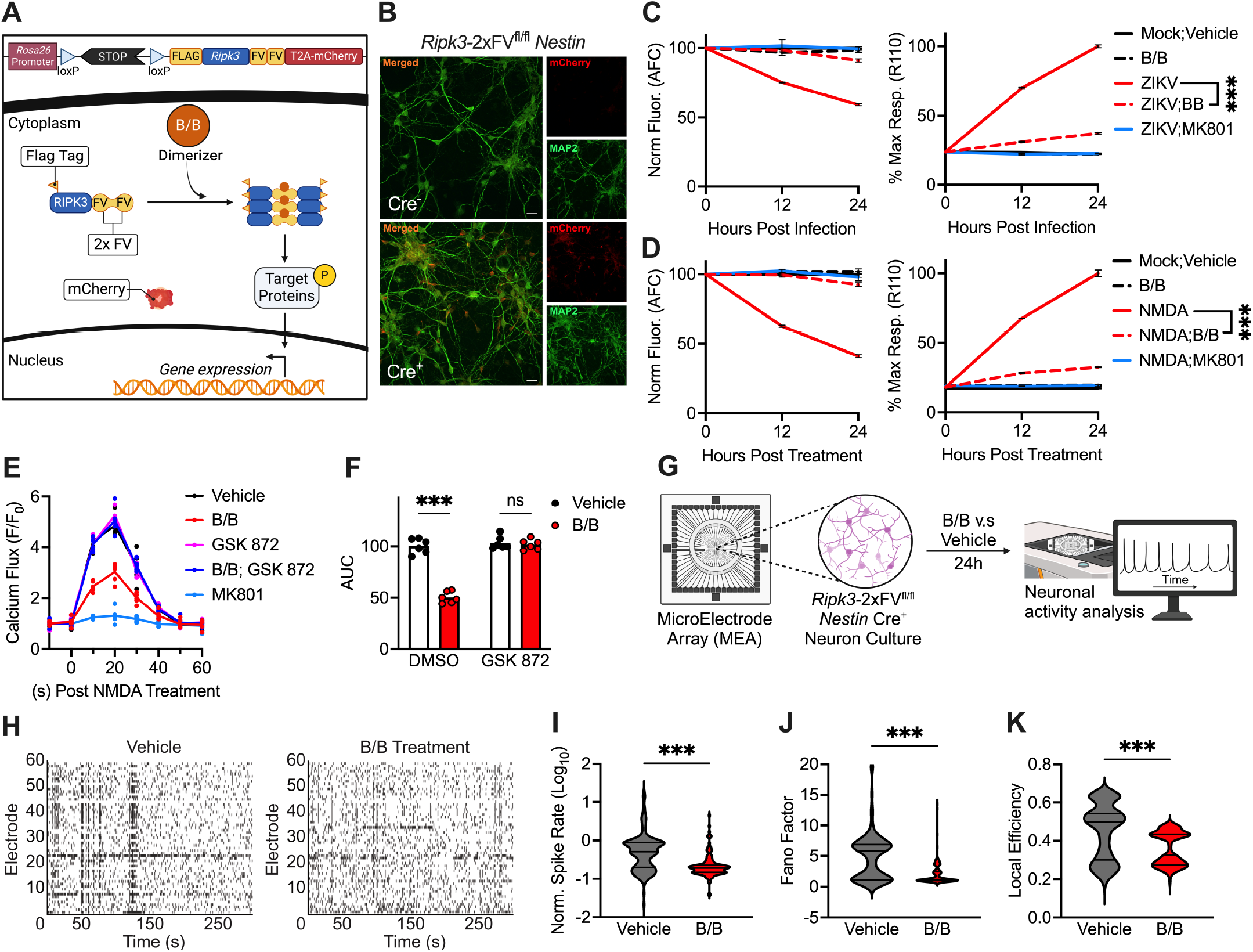
RIPK3 modulates neuronal excitability and promotes neuroprotection during excitotoxicity. (A)Schematic of the *Ripk3*-2xFV^fl/fl^ activation system. Transgene expression is controlled by a lox-STOP-lox element and is coupled with a non-fused mCherry reporter. Treatment with B/B induces FV-mediated dimerization, activating RIPK3 which triggers subsequent protein phosphorylation and gene expression. (B)Immunocytochemistry in primary neuron cultures from *Ripk3*-2xFV^fl/fl^ *Nestin* Cre^+^ and Cre^-^ mice, displaying neuronal MAP2 (green) and mCherry (red) staining. Scale bar: 20 microns. (C-D) MultiTox cell viability assay in *Ripk3*-2xFV^fl/fl^ *Nestin* Cre^+^ neurons pretreated for 24h with B/B, followed by exposure to ZIKV (C) or NMDA (D). Panels report live (AFC) and dead (R110) protease activity, as indicated. N = 6 independent cultures per group/condition. (E-F) NMDA-evoked Ca^2+^ dynamics in cortical neuron cultures following 24h pretreatment with specified drugs, measured at 10-second intervals in the presence of Brilliant Calcium Flex reagent (E). Area under the curve (AUC) analysis comparing indicated groups is shown in (F). (G-K) Primary neurons from *Ripk3*-2xFV^fl/fl^ *Nestin* Cre^+^ embryos were cultured on microelectrode arrays. Following 24h treatment with either B/B or vehicle, neuronal activity on individual arrays was recorded and analyzed (G). Representative raster plots displaying activity across individual electrodes over time are shown (H), with quantitative analysis of normalized spike rates (I), Fano factor (J), and local network efficiency (K). N = 3 arrays per group, with each array containing 59 unique recording electrodes. ***p < 0.001. Error bars represent SEM. **See also Figure S2**.

Given these findings, we next evaluated whether RIPK3 activation could modulate NMDAR-dependent Ca^2+^ flux. We measured intracellular Ca^2+^ dynamics following a pulse stimulation with NMDA in neurons expressing RIPK3-2xFV using a cell permeable fluorometric indicator (Brilliant Calcium). While vehicle treated neurons exhibited robust calcium flux in response to NMDA, this response was significantly diminished in neurons pretreated for 24h with B/B (Figure 3E-F). This suppressive effect of RIPK3 activation could be completely abolished in the presence of a pharmacologic inhibitor of RIPK3 kinase activity (GSK 872), suggesting that RIPK3 kinase function was required for this activity. Importantly, the effect of GSK 872 did not arise due to drug-induced changes in neuronal viability (Figure S2D). Calcium flux in this assay could also be completely abolished in the presence of MK801, confirming that we were measuring NMDAR-dependent activity in this assay. Notably, we observed that a short 2h pretreatment with B/B failed to suppress NMDA-induced calcium flux, suggesting that this effect likely does not occur via immediate RIPK3-mediated regulation of NMDAR function but instead requires engagement of downstream signals that require more time to impact neuronal calcium dynamics (Figure S2E-F). Together, these results indicate that RIPK3 kinase activity is capable of suppressing NMDAR-dependent Ca^2+^ flux in neurons.

To further establish whether RIPK3 activation could directly impact neurotransmission, we cultured *Ripk3*-2xFV^fl/fl^ *Nestin* Cre^+^ neurons on microelectrode arrays (MEAs) and recorded spontaneous network activity following 24h treatment with either B/B or vehicle control solution (Figure 3G). Notably, B/B treated cultures displayed a reduction in spontaneous spiking activity (Figure 3H-I), suggesting a decrease in overall neuronal excitability. Furthermore, spiking activity in B/B treated cultures exhibited less variability over time as quantified by the Fano factor statistic (Figure 3J), indicating an overall decrease in the complexity of spike activity within the neural network. We also observed a decrease in local efficiency of signal transmission in cultures treated with B/B (Figure 3K), consistent with an overall diminished level of signal propagation and network connectivity. Collectively, these findings further support the idea that RIPK3 activation directly modulates neurotransmission by dampening synaptic activity, which may represent a mechanism of neuroprotection under excitotoxic conditions.

### RIPK3 activation modulates neural activity *in vivo* during flavivirus infection

To further explore a potential link between RIPK3 function and neural excitability, we returned to our transcriptomic analysis of primary neurons and performed Ingenuity Pathway Analysis (IPA) to assess how RIPK3 expression influenced the representation of “Disease and Function” terms related to neurotransmission or aberrant neural signaling. We observed enhanced activation scores for many neurologic disease terms in *Ripk3*^−/−^ neurons infected with either ZIKV or WNV (Figure 4A). Notably, terms related to disordered neural activity, such as seizure and epilepsy, were particularly enriched in *Ripk3*^−/−^ neurons following ZIKV infection, while other terms, such as locomotion and movement disorders, were enriched for both viruses. Further analysis revealed hundreds of DEGs related to the seizure and epilepsy-associated IPA terms in *Ripk3*^−/−^ neurons following infection with both viruses (Figure 4B), suggesting that RIPK3 modulates a broad program of genes related to neural excitation in the setting of flavivirus infection.

**Figure 4.**
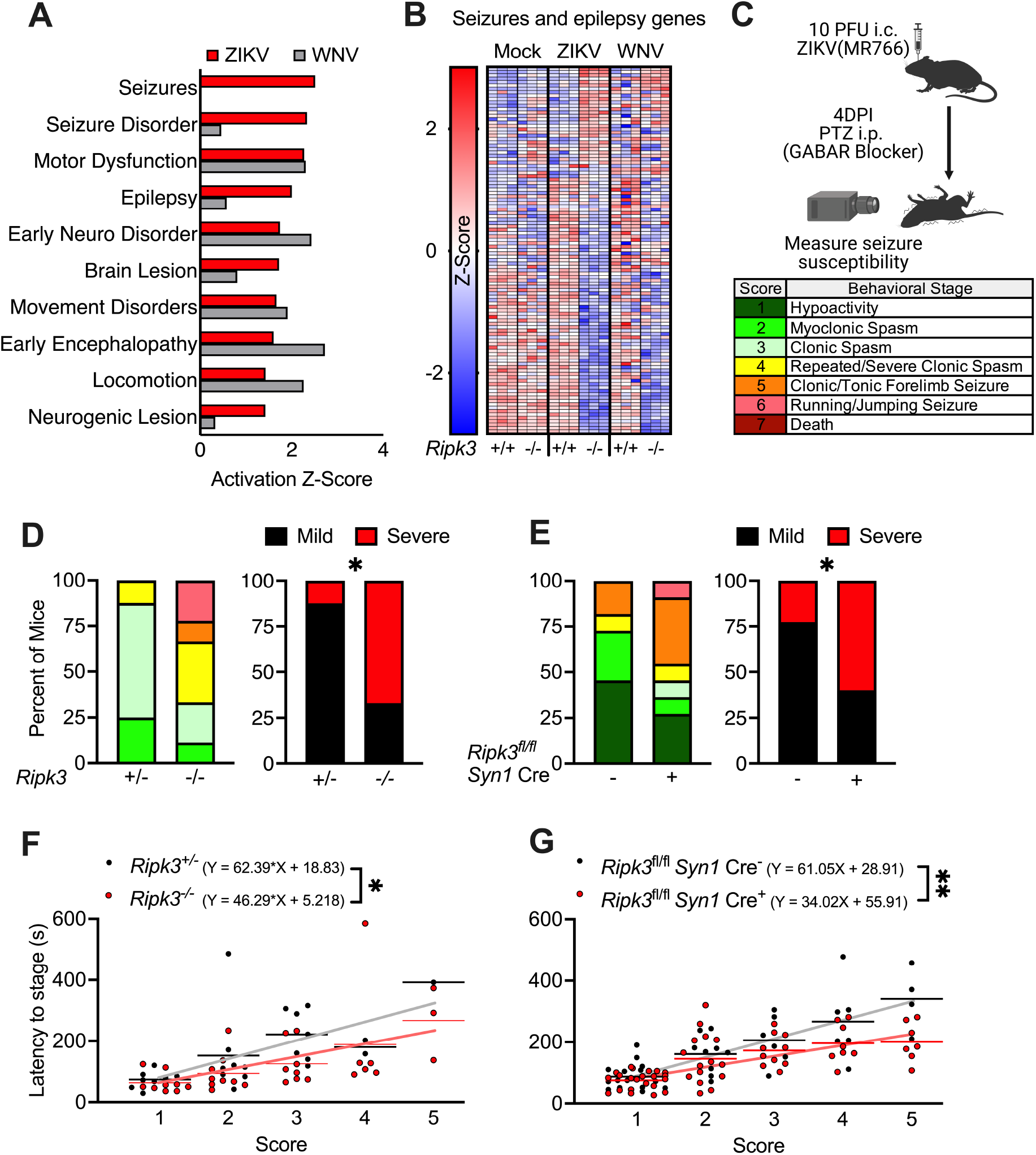
RIPK3 activation modulates neural activity *in vivo* during flavivirus infection. (A)Ingenuity Pathway Analysis of disease and function terms enriched among significant DEGs derived from microarray profiling of primary neurons from specified genotypes after 24h infection with ZIKV or WNV. (B)Heatmap displaying the differential expression of selected genes associated with seizures and epilepsy, derived from microarray profiling of primary cortical neurons from specified genotypes after 24 hours of infection with ZIKV or WNV. (C)Schematic depicting the protocol for inducing seizures with pentylenetetrazol (PTZ) 4 days after intracranial ZIKV infection, including a table explaining the modified Racine Scale of murine seizure stages. (D-E) Proportion of mice reaching indicated behavioral seizure stages, with “severe” seizures defined as stage 4 or higher, across indicated genotypes. N = 8-9 (D) or 15-22 (E) mice per group. (F-G) Latency time in seconds for mice to reach consecutive seizure stages as shown in panels (D) and (E). Linear regression was used to compare overall rates of seizure progression between groups. Latency calculations end at stage 5 as proportionally too few control mice reach higher stages to include in the regression analysis. *p < 0.05, **p < 0.01. Error bars represent SEM. **See also Figure S3**.

We next explored whether RIPK3 influences neuronal activity in the context of flavivirus infection within the intact CNS. To do so, we used an established model of chemically-induced seizure in which mice are injected with pentylenetetrazol (PTZ), a GABA type A receptor antagonist. Following intracranial ZIKV infection, mice were subjected to PTZ administration and seizure severity was recorded using a modified Racine scale^42^ (Figure 4C). We observed that infected *Ripk3*^−/−^ mice demonstrated a significant increase in seizure severity compared to infected littermate controls, both in terms of the overall distribution of scores within genotypes as well as the total proportion of mice reaching a “severe” score defined as 4 or greater on the Racine scale (Figure 4D). Identical experiments conducted on mice harboring neuron-specific *Ripk3* deletion (*Ripk3*^fl/fl^ *Syn1* Cre^+^) revealed similar results (Figure 4E). Both whole-body *Ripk3*^−/−^ and *Ripk3*^fl/fl^ *Syn1* Cre^+^ mice also exhibited reduced latencies in reaching successive stages along the seizure severity scale (Figure 4F-G). Notably, similar experiments revealed no significant differences in PTZ-induced seizure susceptibility between *Mlkl*^−/−^ mice and *Mlkl*^*+/-*^ littermate controls following infection (Figure S3A-S3C), suggesting that RIPK3-mediated suppression of seizure severity was independent of MLKL-dependent mechanisms such as necroptosis, in line with our extensive previous work showing that MLKL is not engaged in the CNS during flavivirus infection^13-15^. Together, these data provide further evidence that RIPK3 activity within the infected CNS can suppress aberrant neural excitation *in vivo*.

### RIPK3 signaling drives activation of the key neural regulatory kinase CaMKII

Given our observation of a neuroprotective function for RIPK3 that is independent of canonical necroptosis signaling, we next sought to define downstream signaling mechanisms by which RIPK3 modulates neuronal excitability. IPA analysis of kinase networks putatively engaged by RIPK3 in primary neurons revealed diminished activation of gene networks regulated by many immunological kinases, including JAK1/2, IRAK4, IKK, and TBK1, in *Ripk3*^−/−^ neurons following both ZIKV and WNV infection (Figure S4A), consistent with established roles for RIPK3 in driving transcription of inflammatory genes. However, we also noted that loss of RIPK3 signaling was associated with diminished activation of genes controlled by CaMKII. This molecule represented an attractive mechanistic candidate in our study, as CaMKII, and the CaMKIIα isoform in particular, is well-established as a modulator of a diverse array of processes related to neurotransmission, including glutamate receptor signaling, long term potentiation, and survival following neurotoxic insults^43,44^. Recent work also suggested that CaMKII may be a substrate of RIPK3 in cardiomyocytes^45^. We therefore questioned whether RIPK3 engages CaMKIIα activation in neurons. To do so, we used mice expressing the chemogenetically activatable RIPK3-2xFV protein under the Synapsin-1 promoter (*Ripk3*-2xFV^fl/fl^ *Syn1* Cre^+^). Importantly, this chimeric RIPK3 protein includes a FLAG tag, facilitating molecular biological analysis of potential binding partners. We treated *Ripk3*-2xFV^fl/fl^ *Syn1* Cre^+^ mice with B/B or vehicle solution for 4h then harvested cerebral cortices and performed protein pulldown of RIPK3-2xFV using beads coated with anti-FLAG antibodies (Figure 5A). Western blot confirmed the expression of the RIPK3-2xFV protein as well as highly efficient pulldown in both treatment groups (Figure 5B and S4B). Notably, chemogenetic activation of RIPK3 induced CaMKIIα activation *in vivo*, as evidenced by detection of phosphorylated (p-Thr286) CaMKIIα (pCaMKIIα) following B/B treatment. Moreover, while p-CaMKIIα was not detected in vehicle treated mice, we observed substantial pulldown of RIPK3-2xFV with both total and phosphorylated CaMKIIα when RIPK3 activation was enforced with B/B, suggesting that CaMKIIα may be a direct substrate of RIPK3 kinase activity, or that the proteins interact indirectly in a complex that ultimately drives CaMKIIα phosphorylation.

**Figure 5.**
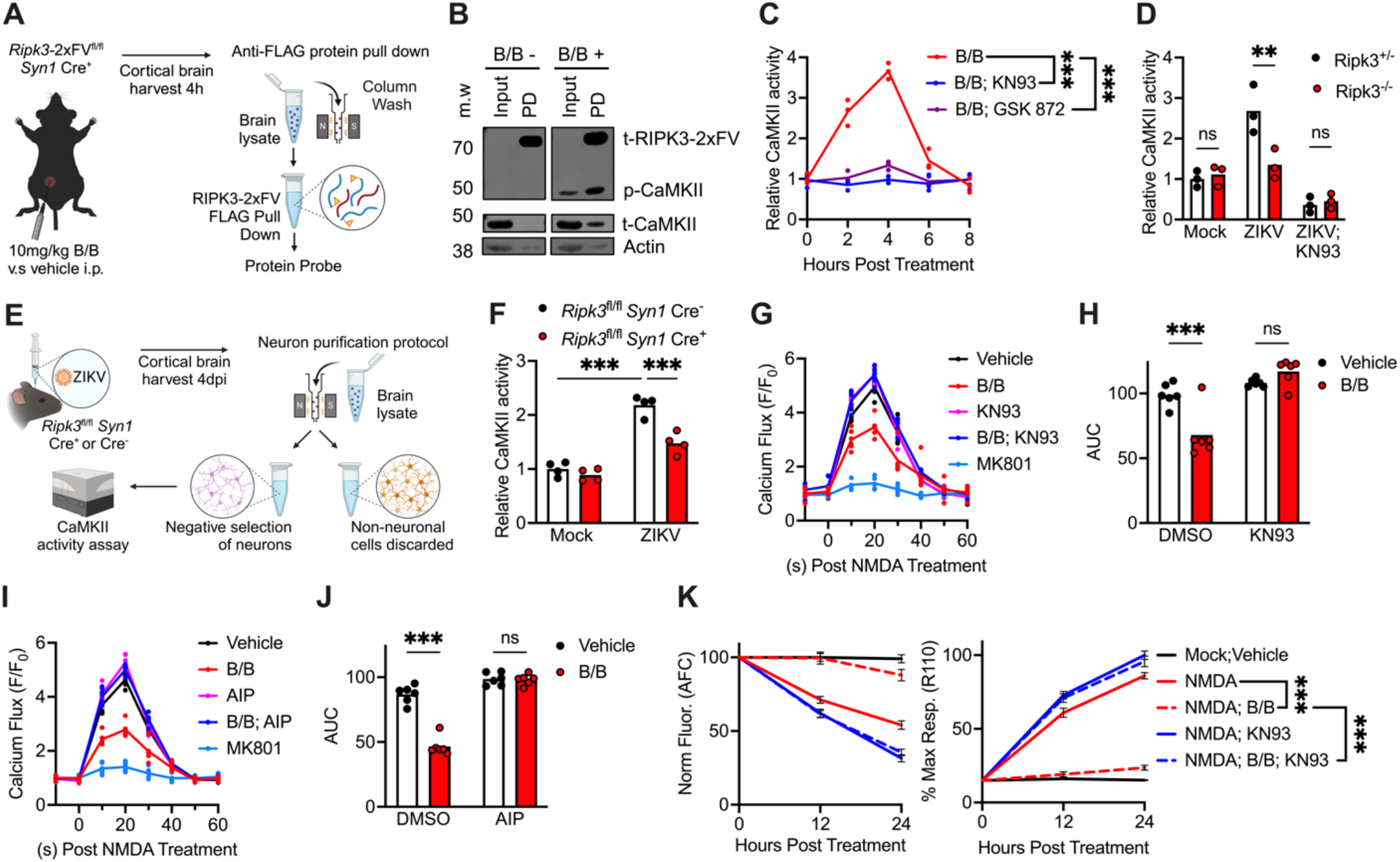
RIPK3 signaling drives activation of the key neural regulatory kinase CaMKII. (A)Schematic illustrating the pulldown assay of FLAG-tagged RIPK3-2xFV protein from cortical brain tissue derived from *Ripk3*-2xFV^fl/fl^ *Syn1* Cre^+^ mice, 4h after i.p. administration of B/B. (B)Western blot analysis of phosphorylated (p)-CaMKII in the input and pulldown (PD) samples, with total (t)-CaMKII, p-CaMKII, and RIPK3 shown for comparison. Actin is used as a loading control. Blots are representative of 3 independent experiments. (C-D) ELISA-based assay of CaMKII activity in *Ripk3*-2xFV^fl/fl^ *Nestin* Cre^+^ primary cortical neuron cultures at indicated time points following treatment with B/B (C) or 24h post-ZIKV infection (D), in the presence of specified inhibitors. (E-F) Schematic of the negative selection MACS protocol used to purify neurons from cerebral cortices of *Ripk3*^fl/fl^ *Syn1* Cre^+^ or Cre^-^ mice 4 days following ZIKV infection (E). (F) shows ELISA-based detection of CaMKII activity in isolated cortical neurons derived from the indicated conditions. (G-J) NMDA-evoked Ca^2+^ dynamics in cortical neuron cultures following 24h pretreatment with specified drugs, measured at 10-second intervals in the presence of Brilliant Calcium Flex reagent (G,I). Area under the curve (AUC) analysis comparing indicated groups is shown in (H,J). (K) MultiTox cell viability assay in *Ripk3*-2xFV^fl/fl^ *Nestin* Cre^+^ neurons treated with B/B homodimerizer for 24h with or without KN93, followed by exposure to NMDA. Panels report live (AFC) and dead (R110) protease activity, as indicated. N = 6 independent cultures per group/condition. **p < 0.01, ***p < 0.001. Error bars represent SEM. **See also Figure S4**.

To further test this idea, we assessed CaMKII activity following sterile RIPK3 activation with B/B in *Ripk3*-2xFV^fl/fl^ *Nestin* Cre^+^ primary neuron cultures using an ELISA-based CaMKII activity assay. We observed a time-dependent increase in CaMKII activity that peaked 4h following B/B administration (Figure 5C). RIPK3-induced CaMKII activation was abolished by cotreatment with a CaMKII inhibitor (KN93) or with an inhibitor of RIPK3 kinase activity (GSK 872), suggesting that RIPK3 kinase activity is required to drive CaMKII activation in this setting. We also observed enhanced CaMKII activity in *Ripk3*^*+/-*^ but not *Ripk3*^−/−^ primary neurons infected with ZIKV, an effect that was also prevented by KN93 treatment, suggesting that RIPK3-dependent induction of CaMKII activity was not an artifact of our chemogenetic system but also occurs using a physiological stimulus that activates endogenous RIPK3 (Figure 5D). We further confirmed this *in vivo* by infecting *Ripk3*^fl/fl^ *Syn1* Cre^+^ mice or Cre^-^ littermate controls with ZIKV and performing negative selection purification of neurons using magnetic activated cell sorting (MACS) (Figure 5E). We observed robust enhancement of CaMKII activity in sorted neurons derived from Cre^-^ animals following infection, and this effect was significantly blunted in neuron-specific *Ripk3* knockout animals (Figure 5F). These data strongly support a role for RIPK3 in driving neuronal CaMKII activation during infection.

We next assessed whether engagement of CaMKII activation was required for RIPK3-dependent modulation of neuronal physiology. We first assessed calcium flux dynamics by treating *Ripk3*-2xFV^fl/fl^ *Nestin* Cre^+^ neuron cultures overnight with B/B in the presence of either of two CaMKII inhibitors, KN93 and a cell permeable version of the CaMKII-specific inhibitory peptide AIP, which were used at concentrations that did not impact neuronal viability (Figure S4C). We observed that inhibition of CaMKII activity with either reagent prevented suppression of calcium flux in B/B treated neurons (Figure 5G-J). We also evaluated NMDA-induced excitotoxicity in *Ripk3*-2xFV^fl/fl^ *Nestin* Cre^+^ neurons pretreated with B/B with or without KN93 using the MultiTox Cytotoxicity Assay. We observed that pharmacologic inhibition of CaMKII abrogated the pro-survival effect of enforced RIPK3 activation in this assay (Figure 5K). Together, these data suggest that CaMKII is required for the neuroprotective functional outcomes of RIPK3 activation in neurons.

### RIPK3/CaMKII signaling promotes neuroprotection through engagement of CREB and de novo transcription

To further characterize the mechanisms by which the RIPK3/CaMKII signaling axis promoted neuroprotection during excitotoxicity, we next sought to understand how this pathway shapes neuronal gene expression. Our transcriptomic profiling suggested a profound role for RIPK3 in regulating expression of genes relevant for neurotransmission and neuronal survival, and the kinetics of many of our functional assays suggested that engagement of transcription and translation was likely required for RIPK3 to exert its neuroprotective effects in our model systems. Notably, CaMKII is known to influence neural physiology in large part through its activation of the transcription factor CREB, a molecule extensively described to be a key regulator of both neurotransmission and neuronal survival^46,47^. Among its many functions, CREB phosphorylation at Ser133 by CaMKII initiates a transcriptional response that promotes neuroprotection subsequent to NMDAR activation^27,48^. We therefore hypothesized that RIPK3-CaMKII signaling culminates in CREB-mediated transcriptional responses. To test this idea, we assessed CREB activation in neurons using an ELISA-based assay in which phosphorylated CREB in sample lysates is captured on plates coated with the cAMP response element DNA sequence, facilitating colorimetric detection. This assay revealed that *Ripk3*-2xFV^fl/fl^ *Nestin* Cre^+^ neuron cultures treated with B/B exhibited significantly enhanced CREB activation (Figure 6A). However, this effect was abrogated in the presence of either GSK 872 or KN93, suggesting that CREB phosphorylation subsequent to RIPK3 activation requires the kinase activities of both RIPK3 and CaMKII. Further experiments showed that ZIKV infection also resulted in enhanced CREB phosphorylation in *Ripk3*^*+/-*^ but not *Ripk3*^−/−^ neuron cultures, and that ZIKV-induced CREB phosphorylation was also prevented in the presence of KN93 (Figure 6B). We similarly observed enhanced CREB phosphorylation in sorted neurons derived from *Ripk3*^fl/fl^ *Syn1* Cre^-^ control mice following ZIKV infection *in vivo*, while this effect was lost in Cre^+^ littermates (Figure 6C-D). These data suggest a signaling pathway in which RIPK3 drives successive signaling events involving CaMKII activation followed by CREB activation in neurons.

**Figure 6.**
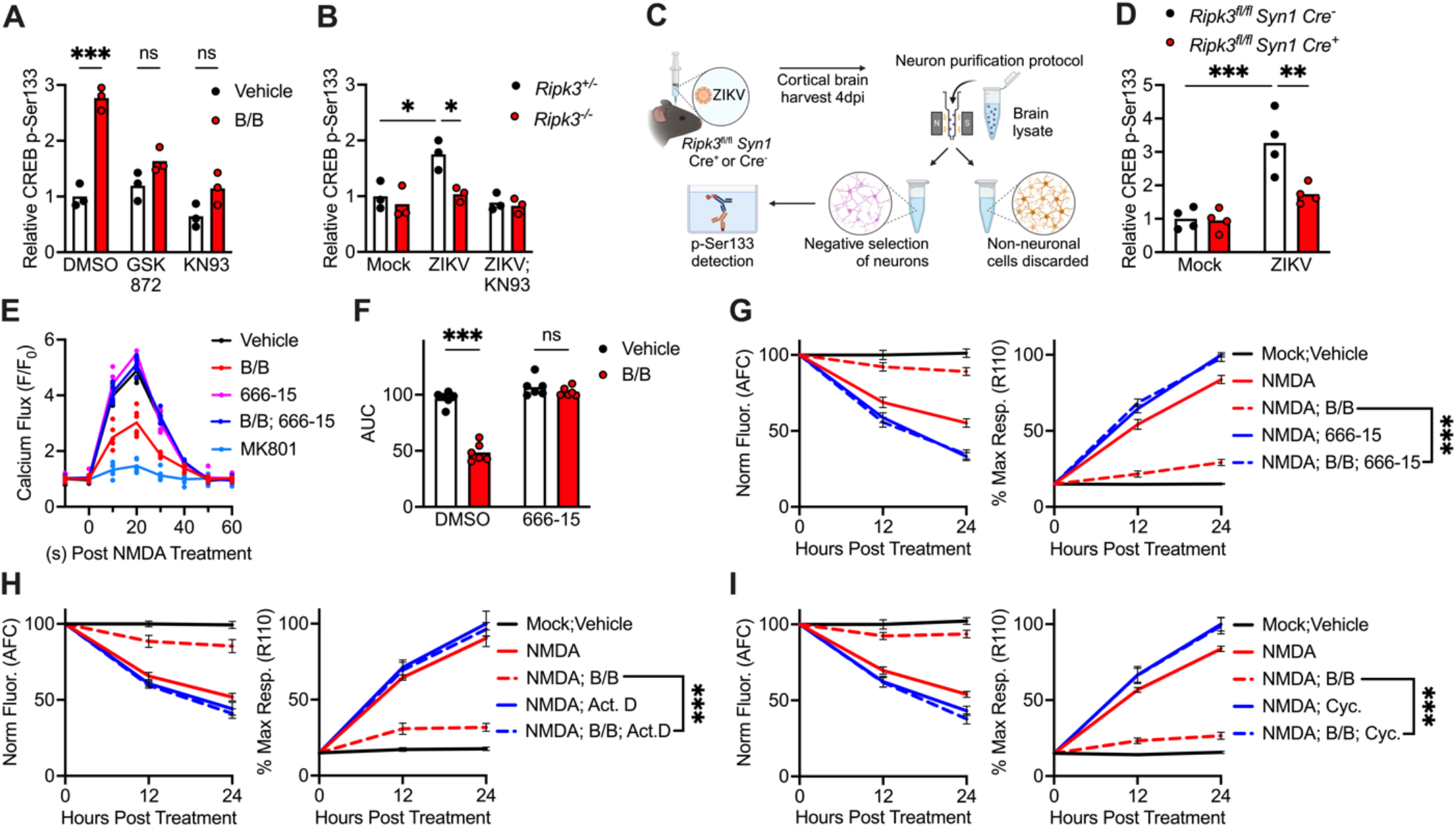
RIPK3/CaMKII signaling promotes neuroprotection through engagement of CREB and de novo transcription. (A-B) Detection of phosphorylated CREB (p-Ser133) in *Ripk3*-2xFV^fl/fl^ *Nestin* Cre^+^ neuron cultures 4h following treatment with B/B and specified inhibitors (A) or 24h following ZIKV infection in indicated genotypes (B). (C-D) Schematic of the negative selection MACS protocol used to purify neurons from cerebral cortices of *Ripk3*^fl/fl^ *Syn1* Cre^+^ or Cre^-^ mice 4 days following ZIKV infection (C). (D) shows ELISA-based detection of phosphorylated CREB in isolated cortical neurons derived from the indicated conditions. (E-F) NMDA-evoked Ca^2+^ dynamics in cortical neuron cultures following 24h pretreatment with specified drugs, measured at 10-second intervals in the presence of Brilliant Calcium Flex reagent (E). Area under the curve (AUC) analysis comparing indicated groups is shown in (F). (G-I) MultiTox cell viability assay in *Ripk3*-2xFV^fl/fl^ *Nestin* Cre^+^ neurons treated with B/B homodimerizer for 24h with or without indicated inhibitors of CREB (G), de novo transcription (H), or de novo translation (I), followed by exposure to NMDA. Panels report live (AFC) and dead (R110) protease activity, as indicated. N = 6 independent cultures per group/condition. **p < 0.01, ***p < 0.001. Error bars represent SEM. **See also Figure S5**.

We next explored the role of CREB in modulating RIPK3/CaMKII-mediated calcium flux after NMDA pulse stimulation. We observed that the CREB inhibitor 666-15 abrogated the suppressive effect of B/B treatment on neuronal calcium flux in response to NMDA, suggesting that CREB-mediated transcription is required for this effect (Figure 6E-F). In contrast, neither of two specific inhibitors of NFκB influenced RIPK3-mediated suppression of neuronal calcium flux (Figure S5A-D). These findings are significant because NFκB has been widely shown to be a central mediator of inflammatory transcription engaged by RIPK3 signaling in a variety of cell types, suggesting that modulation of neurotransmission is likely not a nonspecific effect of inflammatory activation in neurons. We also observed that blockade of CREB activation with 666-15 also prevented the neuroprotective effect of RIPK3 activation in RIPK3-2xFV expressing neuron cultures following exposure to a toxic NMDA stimulus (Figure 6G and S5E). To more firmly establish whether RIPK3-mediated neuroprotection required *de novo* transcription and translation, we performed cell viability experiments in which RIPK3 was chemogenetically activated in the presence of the transcription inhibitor actinomycin D or the translation inhibitor cycloheximide at concentrations that were not intrinsically toxic to cultured neurons (Figure S5F). These experiments showed that inhibition of either process abrogated the ability of RIPK3 activation to suppress NMDA-induced neuronal cell death (Figure 6H-I). Together, these data support a model in which RIPK3 preserves neuronal survival through engagement of CaMKII-dependent activation of CREB, which induces expression of transcriptional targets that dampen neuronal excitability and promote neuroprotection during excitotoxic insults.

### RIPK3 activation induces a CaMKII- and CREB-dependent neuroprotective transcriptional program

To further assess how the RIPK3/CaMKII/CREB pathway influences neuronal transcription, we performed bulk RNA sequencing (RNAseq) of primary neuron cultures expressing RIPK3-2xFV treated for 24h with B/B or ethanol vehicle in the presence of KN93, 666-15, or DMSO vehicle. Chemogenetic activation of RIPK3 robustly altered the neuronal transcriptome, as indicated by 6,313 significant DEGs in our RNAseq analysis (Figure 7A). Notably, B/B treatment impacted a much smaller set of transcripts in the presence of KN93 compared to inhibitor-matched control cultures (1,953 DEGs), suggesting that a large proportion of the RIPK3-dependent transcriptional response was dependent on activation of CaMKII function (Figure 7B). Inhibition of CREB using 666-15 had a comparatively smaller impact compared to blockade of CaMKII, but nevertheless substantially decreased the number of significant DEGs induced by B/B (4,572 DEGs) (Figure 7C). These data suggest that CREB is likely only one of potentially many targets of CaMKII activity downstream of RIPK3 activation in neurons. However, given our functional data demonstrating a requirement for CREB in RIPK3-mediated neuroprotection, we sought to further characterize the CREB-dependent portion of the transcriptomic shift induced by B/B. IPA revealed that B/B treatment altered several pathways with relevance to our study, including terms related to Ca^2+^ influx and abundance, as well as epilepsy and neurotransmitter levels (Figure 7D). In contrast, these transcriptional shifts were notably blunted in the setting of either CaMKII or CREB inhibition and, indeed, RIPK3 activation even activated rather than suppressed genes associated with Ca^2+^ mobilization in the presence of either inhibitor. Further analysis of additional gene modules relevant to our functional studies revealed that RIPK3 activation suppressed representative genes whose expression are induced in an activity-dependent fashion (Figure 7E), as well as genes related to glutamate receptor signaling (Figure 7F), and epilepsy/seizures (Figure 7G). In contrast, expression within these gene modules was markedly higher in the presence of KN93, while 666-15 treatment essentially resulted in intermediate levels of expression that were nevertheless higher than those observed in cultures treated with B/B and no inhibitor. A converse phenotype was observed in gene modules related to the suppression of cell death (Figure 7H) and neuroprotection (Figure 7I), which were strongly upregulated by B/B alone but were less so when either CaMKII or CREB was inhibited. In contrast, expression of the top DEGs associated with inflammatory signaling in our analysis were not impacted by either CaMKII or CREB inhibition (Figure S6A), suggesting that the CaMKII/CREB-mediated arm of RIPK3 signaling is not directly involved in the traditional inflammatory outputs of this pathway. These patterns of expression are all consistent with the idea that RIPK3 activation suppresses genes associated with neuronal excitation and supports expression of genes associated with neuroprotection, in a strongly CaMKII-dependent manner and at least partially in a CREB-dependent manner.

**Figure 7.**
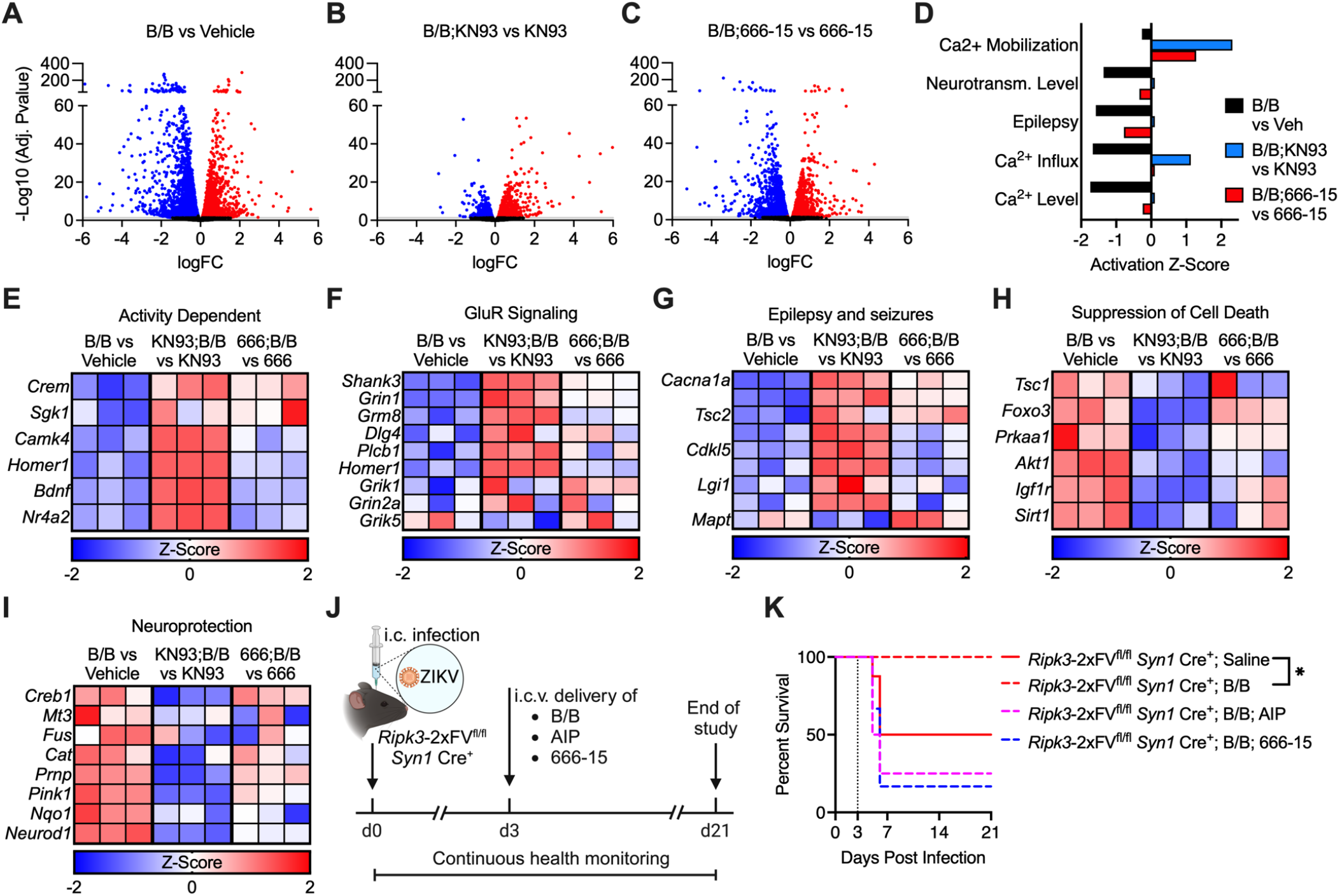
RIPK3 activation induces a CaMKII- and CREB-dependent neuroprotective transcriptional program. (A-I) *Ripk3*-2xFV^fl/fl^ *Nestin* Cre^+^ primary neuron cultures were treated with B/B or ethanol vehicle in the presence of KN93, 666-15, or DMSO vehicle for 24h and then subjected to bulk RNA-sequencing. (A-C) Volcano plots showing differentially expressed genes (DEGs) in indicated comparisons. Data points in red are exhibit upregulated expression, while those in blue exhibit downregulated expression. Genes with an FDR < 0.05 were considered significant. (D) Selected significantly enriched IPA terms showing activation scores within each of the indicated comparisons. (E-I) Heatmaps showing selected significant DEGs associated with indicated pathways. Data are expressed as Z-scores of Log_2_ Fold change values within each comparison. (J)Schematic showing treatment paradigm in which *Ripk3*-2xFV^fl/fl^ *Syn1* Cre^+^ mice were intracranially infected with ZIKV. On day 3 post infection, mice received ICV injection of B/B (or vehicle) combined with AIP, 666-15, or DMSO. Mice were then monitored for survival. (K)Survival analysis of mice in indicated treatment groups. N=5-8 animals/group. *p < 0.05, **p < 0.01. **See also Figure S6**.

Given these results, we performed additional pathogenesis studies to confirm that RIPK3-mediated neuroprotection was dependent on CREB *in vivo. Ripk3*-2xFV^fl/fl^ *Syn1* Cre^+^ mice were infected with ZIKV, followed by intracerebroventricular (i.c.v.) administration of B/B simultaneously with a CaMKII inhibitor (AIP) or CREB inhibitor (666-15) on day 3 post infection (Figure 7J). We observed that chemogenetic activation of neuronal RIPK3 conferred significant protection from ZIKV infection, as B/B treatment resulted in significantly enhanced survival following infection in *Ripk3*-2xFV^fl/fl^ *Syn1* Cre^+^ mice (Figure 7K) but not in Cre^-^ littermate controls (Figure S6B). Strikingly, co-treatment with inhibitors of either CaMKII or CREB significantly abrogated the protective effects of enforced RIPK3 activation in neurons. These results suggest that the CaMKII- and CREB-dependent outputs of RIPK3-mediated transcriptional activation are required to promote host survival during flavivirus encephalitis. They also confirm that, although CREB appears to control only a subset of genes impacted by CaMKII downstream of RIPK3, the CREB-dependent portion of this response is nevertheless critical for the functional outcome of neuroprotection.

## Discussion

Recent work has uncovered complex and nuanced roles for immune signaling in the modulation of neural activity. For example, social behavior has been shown to invoke JAK-STAT-mediated transcription that controls neural activity, suggesting evolutionary links between canonical immune and neurologic signaling pathways^49^. More recently, the molecular basis of memory formation has also been shown to involve innate immune sensing of endogenous genomic damage induced by neural activity^50^. In the context of disease, pathogen sensors and inflammatory cytokines have been implicated in a variety of neurologic and behavioral outcomes^51-55^, including sickness behavior^56-58^, and inflammation generally is known to drive pathologic processes in neurons including aberrant synaptic pruning^6,59,60^, excitotoxicity^23,61,62^, and epileptogenesis^63,64^. Our results describe a nexus of neuroimmune signaling in which activation of RIPK3 dampens neuronal excitability, which in turn promotes survival in the presence of excitotoxic levels of glutamate during CNS viral infection. These results add an additional dimension to the idea of neuroimmune control of neurotransmission by demonstrating a physiological role for this process in host protection and the suppression of viral neuropathogenesis.

Our work also adds new insight to our expanding understanding of the adaptation of innate immunity in neurons, whose unique and vital roles in organismal health necessitate tight regulation of cell fate and survival. Extensive previous work has shown that neurons are resistant to programmed cell death^65^, including RIPK3-mediated necroptosis^10^. We and others have shown that the cell-death independent functions of RIPK3 are critical for host control of CNS viral infection through the well-established functions of this kinase in promoting immunological gene expression^13-15^. Here, our findings underscore an even more extensive role for RIPK3 in controlling neuronal biology during infection, including unexpected functions in regulating excitatory neurotransmission. The potential for the pleiotropic kinase CaMKII^66-68^ to be a direct substrate of RIPK3 in neurons (and other cell types) suggests a host of additional roles for RIPK3 and related proteins in regulating fundamental aspects of cell biology that extend beyond canonical immune and/or cell death processes. Whether these putative non-canonical functions of RIPK3 represent a specific evolutionary adaptation of this pathway in neurons is unknown, and an intriguing alternative hypothesis is that regulation of cell biology via transcriptional control represents the more evolutionarily ancient function of this protein. This original function may be uniquely preserved in neurons (and possibly other postmitotic cell types) but has been obscured by the later development of RIPK3-driven cell death in the necroptosis-susceptible cell types in which this pathway has been traditionally studied.

In either case, the exact mechanisms that protect neurons from necroptosis during RIPK3 activation remain to be discovered. Notably, recent work has shown that overexpression of MLKL in cortical neurons does not render them susceptible to necroptosis following RIPK3 activation^69^, suggesting the existence of unique or at least particularly strong regulators that restrain MLKL-driven cell death in neurons. Others have noted potential mechanisms that may underlie neuronal resistance to necroptosis, such as ESCRT-III-mediated exocytosis of MLKL and membrane repair^69,70^. Our results showing strong engagement of CREB following RIPK3 activation provide an additional possibility, particularly given the well-established roles for CREB target genes in preserving neuronal viability across a broad variety of disease states and insults^46,71-73^. Notably, CREB is known to suppress apoptosis through several mechanisms, such as upregulation of pro-survival BCL-2 family proteins^74,75^, and thus may actively suppress necroptosis through transcriptional control of programmed cell death regulatory proteins. CREB has also been shown to suppress the inflammatory activity of NF-κB through various mechanisms, including competition for the coactivator CREB-binding protein (CBP)/p300, which is used for transcriptional activation by both CREB and NF-κB^76,77^. As NF-κB is intricately linked with both RIPK signaling and cell survival, this connection may also provide clues concerning the unique properties of RIPK3 signaling in cell types such as neurons in which CREB is particularly active.

Finally, our findings also provide new insight into the central role of aberrant neural excitation in the pathogenesis of viral encephalitis. While RIPK3 signaling appears to represent an endogenous mechanism of neuroprotection during flavivirus-induced neurotoxicity, this phenomenon supports the idea that pharmacologic interventions that suppress glutamatergic neurotransmission and/or engage CREB-dependent neuroprotection may have therapeutic potential during neuroinvasive flavivirus infection, particularly during acute encephalitis^23^. The role of hyperexcitation in neuronal cell death during these infections also raises the possibility that non-pharmacologic strategies such as neuromodulation^78,79^ may have some benefit, though testing this hypothesis will require extensive mechanistic study. In any case, identifying new strategies to preserve CNS health during flavivirus infections is of critical importance given the significant and growing burden flaviviruses pose to global public health. Flaviviruses, including WNV and ZIKV, continue to circulate on multiple continents and are likely to continue causing recurrent epidemics in coming years^80,81^. Moreover, climate change continues to increase the endemic range of the mosquito vectors that transmit several medically significant flaviviruses to areas where they are not currently present^82^. As numerous other emerging flaviviruses (Tick-borne encephalitis virus, Spondweni virus, Usutu virus, others) are both neurovirulent and possess epidemic potential^83^, continued work aimed towards understanding mechanisms of neuropathogenesis and neuroprotection during flavivirus infection is particularly urgent.

## Acknowledgements

This work was supported by R21 NS130282 (to BPD and BLF) and R01 NS120895-S2 (to BPD and IE). IE was supported by an HHMI Gilliam Fellowship.

## Author Contributions

Conceptualization: IE, BLF, BPD; Investigation: IE, BDB, NP, ML, TWC, MM, BJV, CA, BPD; Analysis: IE, BDB, NP, TWC, MM, BJV, BPD; Resources: BLF, BPD; Writing – Original Draft: IE, BPD; Writing – Review and Editing: IE, BJV, BLF, BPD; Supervision: CA, BLF, BPD; Funding Acquisition: BLF, BPD.

## Methods

### Mouse lines

*Ripk3*^−/−^, *Mlkl*^−/−^, *Ripk3*^*fl/fl*^, *Ripk3-2xFV*^*fl/fl*^, *Syn1-*Cre, *Nestin-*Cre mice in this study were bred and housed under specific-pathogen free conditions in Nelson Biological Laboratories at Rutgers University. All lines were congenic to the C57BL/6J background (Jackson Laboratories Strain 000664). Primer sequences used to genotype all transgenic mouse lines are listed in Table S1. PCR was conducted using Platinum Taq DirectPCR and Lysis buffer (Invitrogen) using genomic DNA extracted from ear tissue following standard procedures. All mouse studies were performed in 8-12 week old animals of both sexes, following protocols approved by the Rutgers University Institutional Animal Care and Use Committee (IACUC).

### Viruses and virologic assays

ZIKV strain MR766 was provided by the World Reference Center for Emerging Viruses and Arboviruses (WRCEVA). WNV strain WN02-Bird 114^84^ was generously provided by Dr. Bobby Brooke Hererra (Rutgers Robert Wood Johnson Medical School). Vero cells (ATCC #CCL-81) were used to propagate the virus and maintain viral stocks. These cells were cultured in DMEM (Corning #10-013-CV) enriched with 10% heat-inactivated fetal bovine serum (FBS) (Gemini Biosciences #100-106), 1% penicillin-streptomycin-glutamine (Gemini Biosciences #400-110), 1% amphotericin B (Gemini Biosciences #400-104), 1% non-essential amino acids (Cytiva #SH30238.01), and 1% HEPES buffer (Cytiva SH30237.01).

Viral titers were quantified using a plaque assay on the cultured Vero cells. The basal medium for the assay was 1X EMEM (Lonza #12-684F), supplemented with 2% heat-inactivated FBS, 1% penicillin-streptomycin-glutamine, 1% amphotericin B, 1% non-essential amino acids, 1% HEPES, 0.75% sodium bicarbonate (VWR #BDH9280), and 0.5% methyl cellulose (VWR #K390) to form the overlay medium. At 4 days post-infection, the overlay media was removed and cells were fixed/stained using 10% neutral buffered formalin (VWR #89370) and 0.25% crystal violet (VWR #0528) to visualize and count plaques.

### Cell culture and infection

Primary cerebral cortical neurons were derived from E15 mouse embryos as previously described^15^. Neural tissue was harvested and processed using a Neural Tissue Dissociation Kit-T (Miltenyi, #130-093-231) using both male and female embryos. Neurons were seeded onto multi-well plates that had been pre-coated with 50μg/mL Poly-D-Lysine (Thermo Fisher Scientific, #A3890401). Cultures were sustained in Neurobasal-Plus Medium (Thermo Fisher Scientific, #A3582901), enhanced with B-27-Plus supplement (Thermo Fisher Scientific, #A3582801). All culture experiments were performed at 21 days in vitro (DIV). For ZIKV infection experiments, neuronal cultures were infected at a multiplicity of infection (MOI) of 0.1.

### Transcriptomic analysis

Transcriptomic analysis was performed using two distinct data sets. The first was a microarray analysis previously published by us and others using primary cortical neuron cultures derived from both *Ripk3*^*+/+*^ and *Ripk3*^−/−^ mice infected with ZIKV-MR766 or WNV-TX for 24 hours (GEO accession number: GSE122121). The second was a new analysis derived from bulk-RNA sequencing of primary cortical neurons expressing RIPK3-2xFV. Neurons were treated with B/B or vehicle solution in the presence of KN93, 666-15, or control solution for 24 hours prior to harvest. Library preparation and Next Generation Sequencing was performed by Azenta Life Sciences (Piscataway, NJ). RNA yield and sample quality were assessed using Qubit (Invitrogen) and TapeStation (Agilent). Sequencing was performed on the Illumina HiSeq platform using 2 x 150-bp paired-end reads. Sequence reads were cleaned of adapters and low-quality sections, deduplicated, and aligned to the mouse reference genome via the STAR aligner. Gene expression quantification was carried out by counting unique sequences within gene exons. Differentially expressed genes were identified using DESeq2. The GEO accession number for this dataset will be available upon publication.

### Mouse infections and tissue harvesting

Intracranial infections were administered by injecting 10 plaque forming units (PFU) of ZIKV in 10 μl of Hank’s Balanced Salt Solution (HBSS) into the brain’s third ventricle. Subcutaneous infections were administered by injecting 10^3^ PFU of WNV in 50μl of HBSS into a rear footpad. At specified time points post-infection, mice underwent perfusion with 30 mL of cold phosphate-buffered saline (PBS) to prepare for subsequent analyses. Tissues were either processed for adult neuron isolation using magnetic activated cell sorting (MACS) (Miltenyi 130-126-602), for protein pulldown with the Miltenyi μMACS FLAG-protein isolation kit (Miltenyi 130-101-591), or preserved in TRI Reagent (Zymo, #R2050-1) for gene expression analysis.

### Cell Death Assays

Cell viability was evaluated using two distinct methods: the MultiTox Cytotoxicity Assay (Promega, G9201) and the ATP-based CellTiter-Glo Assay (Promega, G7573). The MultiTox Cytotoxicity Assay simultaneously measures live cell (AFC) and dead cell (R110) protease activities within each culture well, providing two complementary data points. AFC signal was normalized to group means of mock or vehicle controls within experiments, while R110 signal were normalized as a percentage of the maximum response observed among groups within a given experiment. The CellTiter-Glo Assay determines cellular ATP levels, serving as an indicator of metabolically active cells. The luminescence output, reflective of ATP concentration, was normalized to group mean values for mock or vehicle controls. Both fluorescence and luminescence signals in these assays were quantified using a SpectraMax iD3 plate reader (Molecular Devices).

### Calcium Flux Assay

Calcium flux dynamics in primary cortical neuron cultures were evaluated using the Brilliant Calcium Flex kit (IonBiosciences, 10000), according to manufacturer’s instructions. Briefly, following experimental treatments, neuron cultures were incubated with a cell membrane-permeable fluorogenic calcium sensor for 1h at 37°C. Cells were then stimulated with a pulse of 100μM NMDA and fluorescent signal was recorded at 10-second intervals for a total duration of 60 seconds.

### Chemical reagents

The following chemical reagents were used in cell culture experiments: GYKI-52466 (1μM, Sigma, G119), MK801 (1μM, Sigma, M107), GSK 872 (1μM, Tocris, 6492), B/B Homodimerizer (200nM, Takara, 635058), L-Glutamic acid (100-1,000μM, Sigma, G1251), N-Methyl-D-aspartic acid (20μM-100μM, Sigma, M3262), KN93 (1μM, Sigma, K1385), myristoylated AIP (1μM, Tocris, 5959), 666-15 (1μM, Tocris, 5661), Actinomycin D (10 ng/ml, Sigma, A1410), Cycloheximide (1 μg/ml, Sigma, C7698), JSH-23 (50μM, Selleckchem, S7351), BAY 11–7085 (100μM, Tocris, 1743). For in vivo injections, the following chemicals were used: GYKI-52466 (1μg/g), MK801 (0.06μg/g), B/B Homodimerizer (10μg/g).

### Quantitative real-time PCR

Total RNA was extracted from primary neuron cultures using the Qiagen RNeasy Mini Kit (Qiagen, 74106), adhering to the manufacturer’s protocol. RNA concentration was determined using a Quick Drop device (Molecular Devices). Complementary DNA (cDNA) was then synthesized using the qScript cDNA Synthesis Kit (Quantabio, 95047). Quantitative RT-PCR (qRT-PCR) was performed utilizing SYBR Green Master Mix (Bio-Rad, CA1725125) on a QuantStudio5 instrument (Applied Biosystems). Cycle threshold (CT) values for the genes under study were normalized to the CT values of the housekeeping gene 18S (CT_Target − CT_18S = ΔCT). Data were further normalized to baseline control values (ΔCT_experimental − ΔCT_control = ΔΔCT (DDCT)). A list of primer sequences used in the study is provided in Table S1.

### Immunofluorescence

Fluorescent immunocytochemistry was performed following fixation of cells in 4% paraformaldehyde for 10 minutes, followed by blocking with 10% goat serum for 15 minutes. Cells were probed with primary antibodies, including rabbit anti-mCherry (Rockland, 600-401-P16) and chicken anti-MAP2 (Abcam, ab5392), for 1 hour at room temperature. For secondary detection, goat anti-rabbit 594 (ThermoFisher, A-11012) and goat anti-chicken 488 (ThermoFisher, A-11039) antibodies were applied for 15 minutes at room temperature. Nuclei were stained with DAPI. Cells were gently washed with 1X PBS between each step to remove unbound antibodies. Imaging was carried out using an Airyscan fluorescent confocal microscope (Carl Zeiss, LSM 800).

### Microelectrode array (MEA) recordings

MEAs (Multi Channel Systems, 60MEA200/10iR-Ti) consist of 59 titanium nitride (TiN) working electrodes with a diameter of 10 μm with 200 μm spacing between electrodes and one internal reference electrode. Prior to recording, MEAs were treated with oxygen plasma for 30 seconds using a PX-500 Plasma System (March Instruments), 100 μg/mL of poly-D-lysine overnight (Sigma, P0899), and 10 μg/mL Laminin (Sigma, L2020) for 2h prior to culture of primary neurons. For recording, culture medium was removed and 1 mL of warmed MEA recording solution^85,86^ was added to each MEA. Each culture was allowed to equilibrate with the MEA solution for 5 minutes in the 37°C incubator before recording.

Recordings were conducted with a MEA2100-Lite-System (Multi Channel Systems) that was maintained at 37°C using the TC02 temperature controller (Multi Channel Systems). The Multi Channel Experimenter software (Multi Channel Systems) was used to record the extracellular potential at each electrode for 5 minutes using a sampling rate of 20 kHz. After recording, MEA recording solution was removed and the conditioned cell culture medium was returned to the MEA.

### MEA signal processing and analysis

All signal processing was conducted using MATLAB (Mathworks). All signals were filtered with a fourth-order Butterworth bandpass filter (300 – 3,000 Hz) to remove low frequency signals and a 60 Hz comb filter was used to remove noise generated by the hardware^87^. An adaptive thresholding algorithm was used to detect ‘spikes’ in the extracellular recordings. For each 10 second interval on each working electrode, a threshold was defined as 4.5 standard deviations times the background noise of that time interval. Whenever the magnitude of the signal exceeded that threshold, a spike was recorded. A minimum interspike interval (ISI) of 2 msec ensured that the same spike was not recorded multiple times. Each electrode had a spike rate calculated as the number of recorded spikes in that channel per minute of recording. Fano factor, a measure of the distribution of spikes over the recording time, was calculated by dividing the recording time into 100 msec bins and counting the number of spikes per bin. The Fano factor was then calculated by dividing the variance of spikes per bin by the mean number of spikes per bin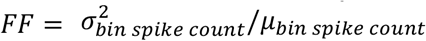. The Brain Connectivity Toolbox (BCT)^88^ was used to characterize network modularity and efficiency. For each electrode, we counted the number of spikes occurring in each 1 msec bin. Functional connectivity matrixes were generated by computing the cross-correlation between the binned spike-trains of each pair of working electrodes using a maximum lag-time of 20 msec. BCT functions for modularity, global efficiency, and local efficiency were then used to quantify those variables.

### Seizure Assay

Seizures were induced by a single i.p. injection of pentylenetetrazole (PTZ) (Sigma-Aldrich, 54-95-5) at a dose of 40μg/g of body weight. Behavioral responses were evaluated using a modified version of the Racine scale as previously described^42^. Adaptation of the scale included broader distinctions between myoclonic and clonic seizures and expanded criteria for generalized hypoactivity. Video recordings of mouse seizures were scored by an operator blinded to the experimental condition of each subject.

### FLAG Pulldown and Western Blot

Pulldown of FLAG-tagged RIPK3-2xFV protein was conducted using the DYKDDDDK Isolation Kit (Miltenyi, 130-101-591) following the manufacturer’s instructions. For Western blot analysis, primary antibodies against the following targets were used: RIPK3 (Cell Signaling Technology, 95702S), CaMKIIα (ThermoFisher, MA1-048), phospho-CaMKIIα (Thermo Fisher Scientific, PA1-4614), and Actin (Sigma-Aldrich, SAB3500350). Secondary antibodies included Goat anti-rabbit IRDye 800CW (Licor Biosciences, 925-32218), Goat anti-mouse IRDye 800CW (Licor Biosciences, 925-32210), and Donkey anti-chicken IRDye 680RD (Licor Biosciences, 926-68075). The immunoblots were visualized using the Odyssey XF Imaging System, which is equipped with two near-infrared lasers.

### Statistical analysis

Statistical analyses were performed using GraphPad Prism 9. Survival experiments were compared via log-rank test. Most other experiments were compared with appropriate parametric tests, including the Student’s t-test (two-tailed) or two-way analysis of variance (ANOVA) with Tukey’s post hoc test to identify significant differences between groups. A p-value of less than 0.05 was deemed to indicate statistical significance. Unless specified otherwise, all data points represent biological replicates consisting of distinct mice or independent cultures derived from distinct mice.

