## Supplemental Material for "RIPK3 promotes neuronal survival by suppressing excitatory neurotransmission during CNS viral infection"

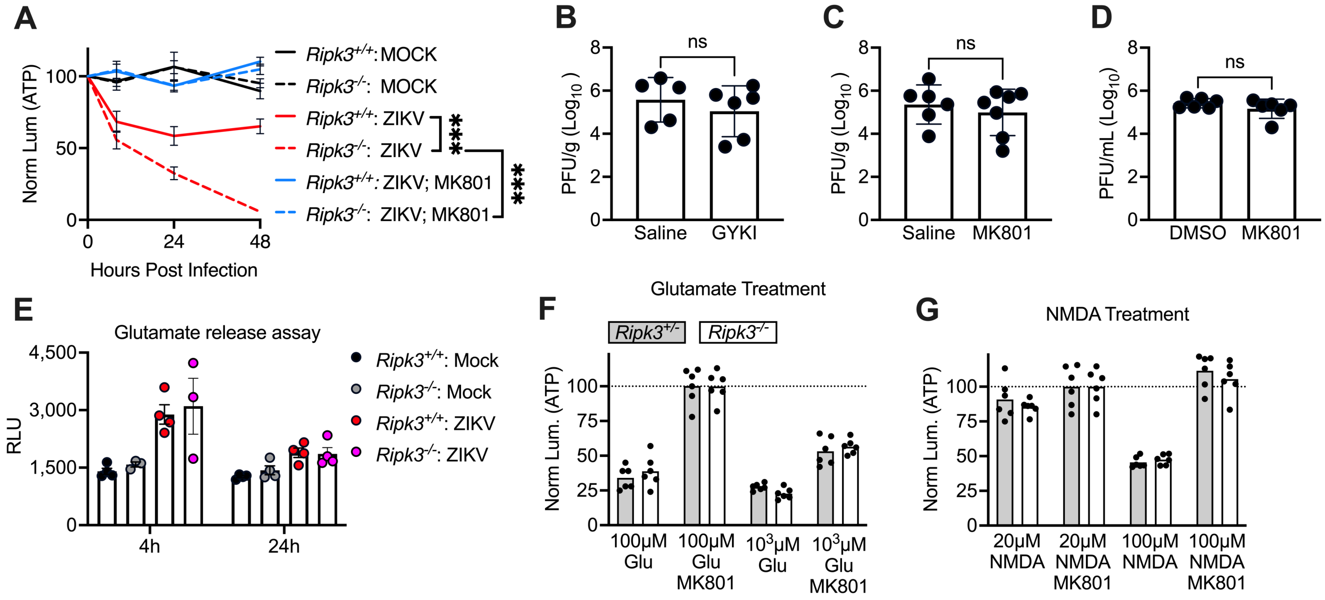


**Figure S1 (Related to Figure 2): Basal effects of GluR inhibitors and *Ripk3*-deficiency on viral burden, viral replication, glutamate release, and excitotoxicity.**
(A) ATP-based viability assay (CellTiter Glo) assessing the impact of glutamatergic signaling on neuronal survival in primary neuron cultures from indicated genotypes following infection with ZIKV. N = 6 independent cultures per group/condition.
(B-C) Viral titer measurements via plaque assay on whole brain homogenates at 4 days post infection from B6/J mice intracranially infected with ZIKV and treated daily (i.p.) with GYKI-52466 (B) or MK801 (C).
(D) Plaque assay measurement of supernatant viral titers in WT neuron cultures pre-treated with MK801 and infected with ZIKV for 24 hours.
(E) Glutamate release assay (Glutamate-Glo) in cortical neuron cultures of indicated genotypes at 4h and 24h following ZIKV infection.
(F-G) ATP-based viability assays to evaluate basal sensitivity to various concentrations of glutamate (F) and NMDA (G) in cortical neuron cultures of specified genotypes, with MK801 used as a control for NMDAR antagonism.

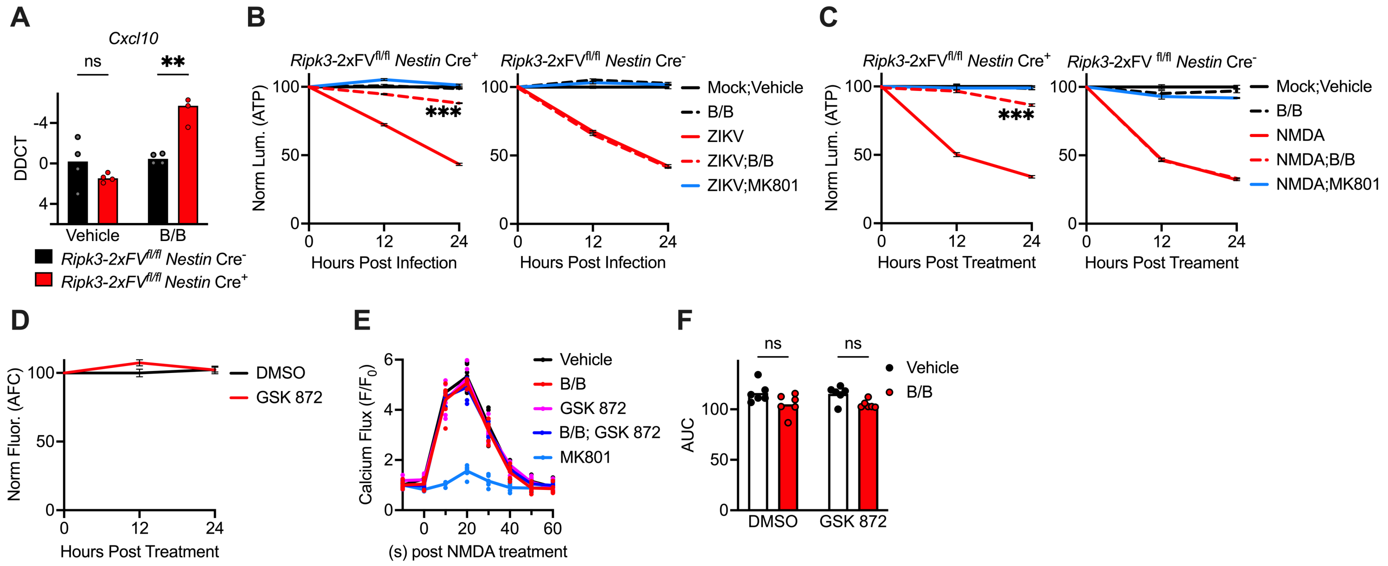


**Figure S2 (Related to Figure 3): Impact of chemogenetic RIPK3 activation on *Cxcl10* expression, neuronal viability, and calcium flux dynamics.**

(A) Measurement of *Cxcl10* transcript levels via qRT-PCR in *Ripk3*-2xFV$^{fl/fl}$ *Nestin* Cre$^{+}$ and Cre$^{-}$ neuron cultures after 24h of B/B treatment.

(B-C) ATP-based viability assays (CellTiter Glo) assessing the impact of 24h B/B pretreatment on primary neuron cultures of indicated genotypes post-ZIKV infection (B) or NMDA exposure (C). N = 6 independent cultures per group/condition.

(D) MultiTox cell viability assay evaluating the effects of the RIPK3 inhibitor GSK 872 on basal neuron viability in WT neuron cultures, measured using live cell protease activity (AFC). N = 6 independent cultures per group/condition.

(E-F) NMDA-evoked $Ca^{2+}$ dynamics in cortical neuron cultures following 2h pretreatment with specified drugs, measured at 10-second intervals in the presence of Brilliant Calcium Flex reagent (E). Area under the curve (AUC) analysis comparing indicated groups is shown in (F).

**p < 0.01, ***p < 0.001. Error bars represent SEM.

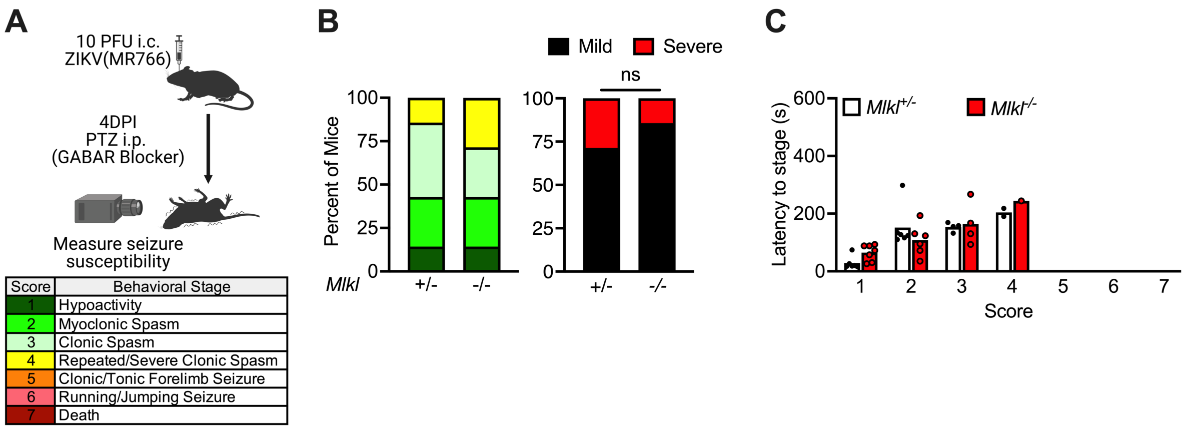


**Figure S3 (Related to Figure 4): MLKL does not modulate neural activity *in vivo* during flavivirus infection.**

(A) Schematic depicting the protocol for inducing seizures with pentylenetetrazol (PTZ) 4 days after intracranial ZIKV infection, including a table explaining the modified Racine Scale of murine seizure stages.

(B) Proportion of mice reaching indicated behavioral seizure stages, with “severe” seizures defined as stage 4 or higher, across indicated genotypes. N = 6-7 mice/group.

(C) Latency time in seconds for mice to reach consecutive seizure stages as shown in panel (B)

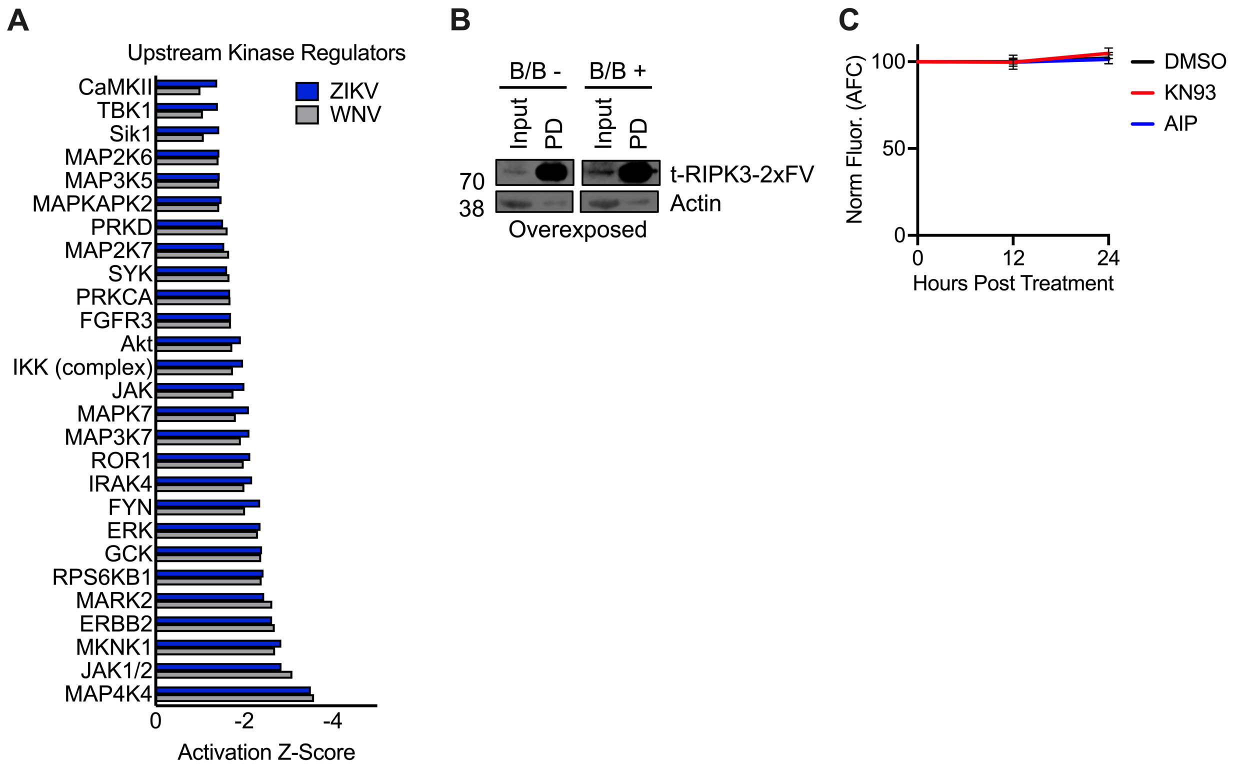


**Figure S4 (Related to Figure 5): RIPK3-mediated regulation of kinase networks during neuronal flavivirus infection and confirmation of RIPK3-2xFV expression in pulldown assays.**

(A) Ingenuity Pathway Analysis comparing kinase network activation in *Ripk3*$^{-/-}$ versus *Ripk3*$^{+/+}$ neurons infected with ZIKV or WNV. Alterations are quantified using activation z-scores.

(B) Overexposure of western blot shown in Figure 5B confirming RIPK3-2xFV expression in input samples. Actin is used as a loading control.

(C) MultiTox cell viability assay evaluating the effects of the CaMKII inhibitors KN93 and AIP on basal neuron viability in WT neuron cultures, measured using live cell protease activity (AFC). N = 6 independent cultures per group/condition.

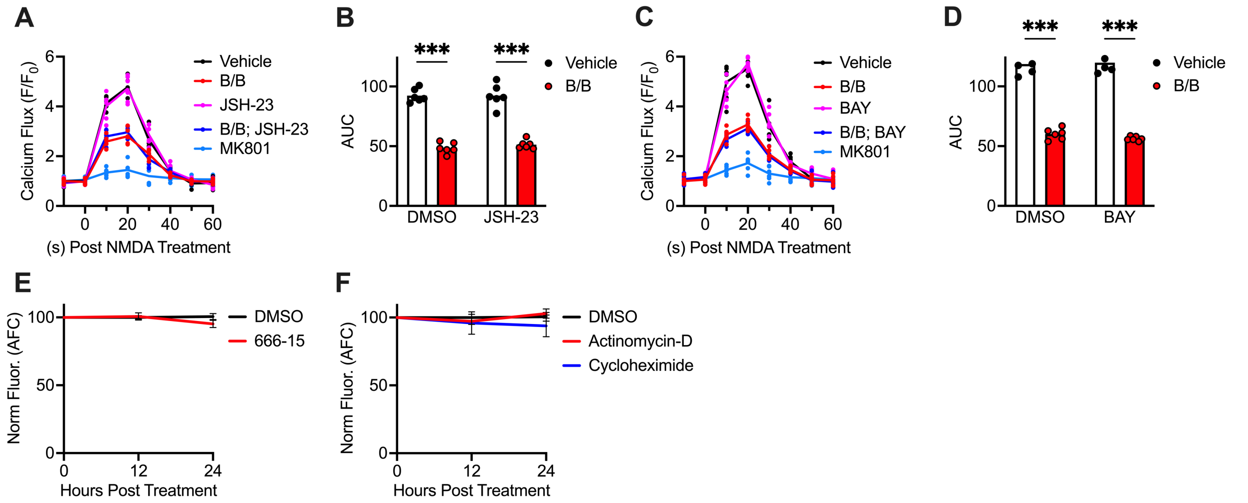


**Figure S5 (Related to Figure 6): The neuroprotective effects of RIPK3 activation do not require NFκB but do require de novo transcription and translation.**
(A-D) NMDA-evoked $Ca^{2+}$ dynamics in cortical neuron cultures following 24h pretreatment with B/B and specified inhibitors of NFκB, measured at 10-second intervals in the presence of Brilliant Calcium Flex reagent (A,C). Area under the curve (AUC) analysis comparing indicated groups is shown in (B, D).
(E-F) MultiTox cell viability assay evaluating the effects of the indicated inhibitors of CREB (E) or transcription/translation (F) on basal neuron viability in WT neuron cultures, measured using live cell protease activity (AFC). N = 6 independent cultures per group/condition.
***$p < 0.001$. Error bars represent SEM.

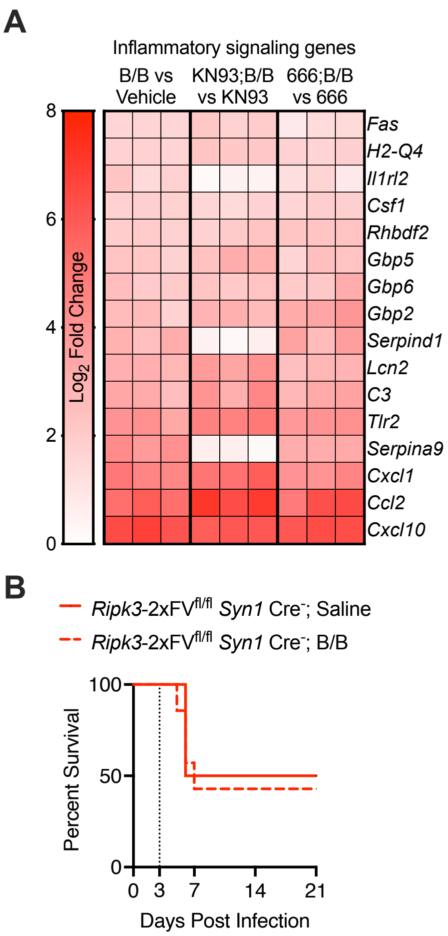


**Figure S6 (Related to Figure 7): CaMKII and CREB are not required for RIPK3-dependent induction of inflammatory genes in neurons.**

(A) *Ripk3*-2xFV$^{fl/fl}$ *Nestin* Cre$^{+}$ primary neuron cultures were treated with B/B or ethanol vehicle in the presence of KN93, 666-15, or DMSO vehicle for 24h and then subjected to bulk RNA-sequencing. Heatmap shows selected significant DEGs associated with inflammatory signaling. Data are expressed as $Log_2$ fold change values within each comparison.

(B) Survival analysis of mice in indicated treatment groups. All mice are Cre$^{-}$ littermate controls of animals used in Figure 7J. N=6 animals/group.

**Table S1: Genotyping and qRT-PCR primer list and description.**

| Target | Note | Primer Sequence (5'-3') | Product Size |
|---|---|---|---|
| Genotyping | | | |
| Ripk3-/- | RIP3_001 | CGCTTTAGAAGCCTTCAGGTTGAC | WT 733bp KO 485 bp |
| | RIP3_002 | GCAGGCTCTGGTGACAAGATTCATGG | |
| | RIP3_003 | CCAGAGGCCACTTGTGTAGCG | |
| Ripk3fl/fl | R3FL_001 | ACGATGTCTTCTGTCAAGTTATG | WT 300bp LoxP 334bp |
| | R3FL_002 | CAGTTCTTCACGGCTCAC | |
| | R3FL_003 | TCTGGTAAGGAGGGTCAC | |
| Ripk3-2xFVfl/fl | ROSA forward | AGCACTTGCTCTCCCAAAGTC | 346bp |
| | ROSA reverse | CCGACAAAACCGAAAATCTGTGGG | |
| | Transgene forward | CGCTTTAGAAGCCTTCAGGTTGAC | 349bp |
| | Transgene reverse | GCAGGCTCTGGTGACAAGATTCATGG | |
| Syn1 Cre | Transgene forward | CTCAGCGCTGCCTCAGTCT | 300bp |
| | Transgene reverse | GCATCGACCGGTAATGCA | |
| | Internal control forward | CTAGGCCACAGAATTGAAAGATCT | 324bp |
| | Internal control reverse | GTAGGTGGAAATTCTAGCATCATCC | |
| Nestin Cre | WT forward | TTGCTAAAGCGCTACATAGGA | WT 246bp Cre 150bp |
| | Common reverse | GCCTTATTGTGGAAGGACTG | |
| | Transgene forward | CCTTCCTGAAGCAGTAGAGCA | |
| Mlkl-/- | MLKL_001 | TATGACCATGGCAACTCACG | WT 498bp KO 158bp |
| | MLKL_002 | ACCATCTCCCCAAACTGTGA | |
| | MLKL_003 | TCCTTCCAGCACCTCGTAAT | |
| qRT-PCR | | | |
| CXCL10 | Forward | GCCGTCATTTTCTGCCTCA | |
| | Reverse | CGTCCTTGCGAGAGGGATC | |
